# A pluripotent stem cell atlas of multilineage differentiation reveals *TMEM88* as a developmental regulator of mammalian blood pressure

**DOI:** 10.1101/2022.10.12.511862

**Authors:** Sophie Shen, Tessa Werner, Samuel W. Lukowski, Stacey Andersen, Yuliangzi Sun, Woo Jun Shim, Dalia Mizikovsky, Sakurako Kobayashi, Jennifer Outhwaite, Han Sheng Chiu, Xiaoli Chen, Gavin Chapman, Ella M. M. A. Martin, Di Xia, Duy Pham, Zezhuo Su, Daniel Kim, Pengyi Yang, Men Chee Tan, Enakshi Sinniah, Qiongyi Zhao, Sumedha Negi, Meredith A. Redd, Joseph E. Powell, Sally L. Dunwoodie, Patrick P. L. Tam, Mikael Bodén, Joshua W. K. Ho, Quan Nguyen, Nathan J. Palpant

## Abstract

Pluripotent stem cells provide a scalable approach to analyse molecular regulation of cell differentiation across multiple developmental lineage trajectories. In this study, we engineered barcoded iPSCs to generate an atlas of multilineage differentiation from pluripotency, encompassing a time-course of WNT-induced differentiation perturbed using modulators of WNT, BMP, and VEGF signalling. Computational mapping of *in vitro* cell types to *in vivo* developmental lineages revealed a diversity of iPSC-derived cell types comprising mesendoderm lineage cell types including lateral plate and paraxial mesoderm, neural crest, and primitive gut. Coupling this atlas of *in vitro* differentiation with Summary data-based Mendelian Randomisation analysis of human complex traits, we identify the WNT-inhibitor protein *TMEM88* as a putative regulator of mesendodermal cell types governing development of diverse cardiovascular and anthropometric traits. Using genetic loss of function models, we show that *TMEM88* is required for differentiation of diverse endoderm and mesoderm cell lineages *in vitro* and that *TMEM88* knockout *in vivo* results in a significant dysregulation of arterial blood pressure. This study provides an atlas of multilineage iPSC differentiation coupled with new molecular, computational, and statistical genetic tools to dissect genetic determinants of mammalian developmental physiology.

## INTRODUCTION

Human pluripotent stem cell (hPSC) models are ideally suited to deconstruct early human developmental processes in a scalable and cell type-specific manner. Significant investment and sector growth in stem cell technologies highlight their importance in developmental biology research, industry drug discovery pipelines, disease modelling, and regenerative medicine^1^. In contrast to *in vivo* cell atlases covering numerous developmental stages across diverse species^2–4^, atlases of *in vitro* differentiation are not widely available. As a consequence, mechanisms governing cell differentiation and fate specification across cell lineages remain inadequately understood^5–7^, creating barriers in guiding the differentiation of hPSCs into cell types to study the mechanism of cell fate decisions^8–10^. Strategies to perturb and map sequential stages of multi-lineage cell differentiation decisions are needed to understand and optimise methods for deriving cell types that model and inform mechanisms of development and disease *in vivo*.

This study presents a multi-scale genetic analysis workflow across *in vitro* and *in vivo* biology to accelerate knowledge gain into mechanisms of development. We first develop a multiplexing method for single-cell RNA sequencing (scRNA-seq), enabled by genomic integration of transcribed barcodes in human induced pluripotent stem cells (iPSCs). Using barcoded iPSCs, we perform a multiplexed perturbation study to characterise the diversification of cell types during multi-lineage differentiation from pluripotency *in vitro*. We evaluate this *in vitro* differentiation atlas using a suite of computational methods to extract the temporal and developmental regulatory mechanisms of differentiating cell lineages.

Translating insights from *in vitro* iPSCs into physiological mechanisms of development or disease remains a significant challenge because *in vitro* models lack complex tissue and organismal phenotypes. We address this limitation by drawing on human complex trait genetic data. Genome-wide association studies (GWAS) and databases mapping genetic diversity to transcriptional variation across tissues (GTEx) reveal how genetic changes influence phenotypic variation in humans. By comparing genetic regulators of *in vitro* cell diversification with GWAS data from UK Biobank, we identify and functionally validate the transmembrane protein, *TMEM88*, as a genetic regulator of cardiovascular blood pressure. Collectively, this study provides new cell-based experimental tools, data resources, and computational workflows that leverage the power of atlas-level multilineage *in vitro* cell diversification to discover genetic determinants of developmental physiology.

## RESULTS

### Derivation of barcoded iPSCs for multiplexed single-cell RNA-seq

Cell barcoding strategies enable sample multiplexing and economise the effort and costs of data generation by scRNA-seq analysis. To enhance sample multiplexing capabilities in human induced pluripotent stem cells (hiPSCs), we designed a cell barcoding method to enable multiplexed single-cell analysis of isogenic human cell lines (**Figure 1a**). 10,000 15-base pair barcodes were randomly generated using an even 25% probability for the presence of each A, C, T and G nucleotides. After excluding barcodes starting or ending with a stop codon or containing repetitive runs of 4 or more nucleotides, we selected 18 barcodes, requiring a minimum Hamming distance of 5 nucleotides between each pair of barcodes (**Table 1**). Using this design, we built barcode plasmids into the AAVS1-2A-Puro gene targeting cassette^11,12^ to enable expression of a barcoded GFP transcript downstream of the ubiquitously expressed CAG at the human *AAVS1* locus after CRISPR-Cas9 integration (**Supplementary File 1**). Transcribed barcodes can thus be captured by any single-cell RNA-seq protocol and amplified in a separate sequencing library to allow each read from pooled experimental samples to be assigned back to its sample of origin without the need for exogenous labelling before sample pooling (**Figure 1a**). With thousands of potential barcode combinations, the extent of multiplexing is only limited by the scale of cell capture technologies. We engineered 18 isogenic hiPSC lines with unique barcodes, each of which passed quality testing for karyotype, pluripotency, and purity (**Figure 1b-e**).

**Figure 1:**
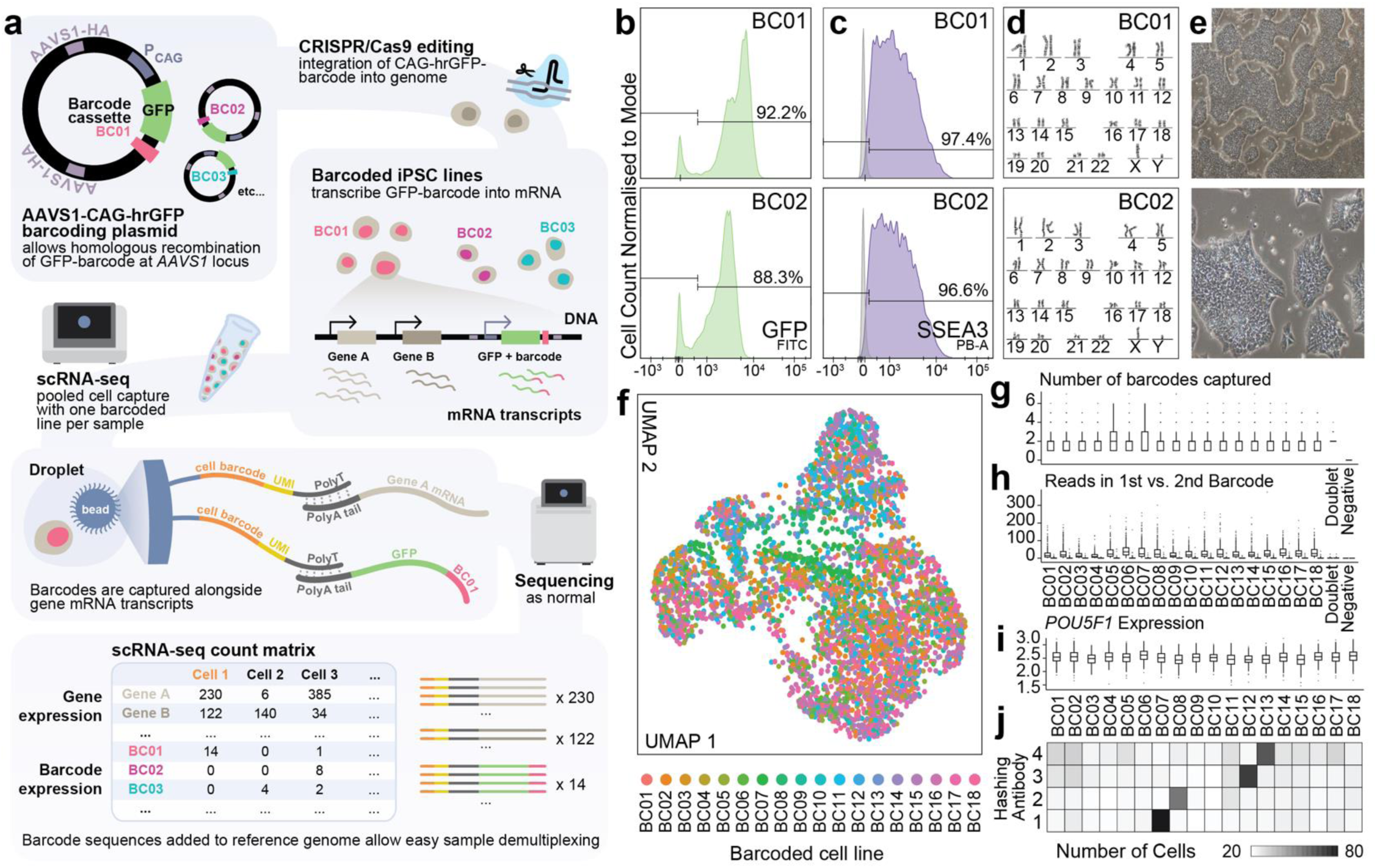
Engineered barcoding of human pluripotent stem cells. **a)** Workflow for CRISPR/Cas9 engineering of barcoded hiPS cell lines to facilitate sample multiplexing of isogenic samples in scRNA-seq experiments. AAVS1-HA: *AAVS1* Homology Arm; hrGFP: humanised Renilla Green Fluorescent Protein; P_CAG_: CAG promoter. See also **Methods** and **Supplementary File 1**. **b-e)** Quality control for barcoded cell lines including representative FACS plots for GPF+ (vector integration) **(b)** and SSEA3+ (pluripotent) cells **(c)**, karyotype **(d)**, and images of pluripotent cell morphology **(e)** for barcoded cell lines BC01 and BC02. **f)** UMAP of pilot scRNA-seq experiment capturing all 18 barcoded cell lines as undifferentiated iPSCs. Points are coloured by the barcode with the highest number of mapped reads per cell. **g)** Number of unique expressed genomic barcodes (out of total 18 possible barcodes) captured per cell in the scRNA-seq experiment grouped by dominant barcode in each cell. See also **Figure S1a**. **h)** Number of reads mapped to the top two most highly mapped genomic barcode in each cell, grouped by dominant barcode in each cell. See also **Figure S1a**. **i)** Expression level of pluripotency gene *POU5F1* (OCT4) in each barcoded line. See also *SOX2* in **Figure S1a**. **j)** Independent validation of barcoding by analysing barcoded cell lines co-labelled with external Cell Hashing antibodies. Antibodies 1-4 were stained to cells in barcoding lines BC07, BC08, BC12, and BC13 respectively.

**Table 1:**
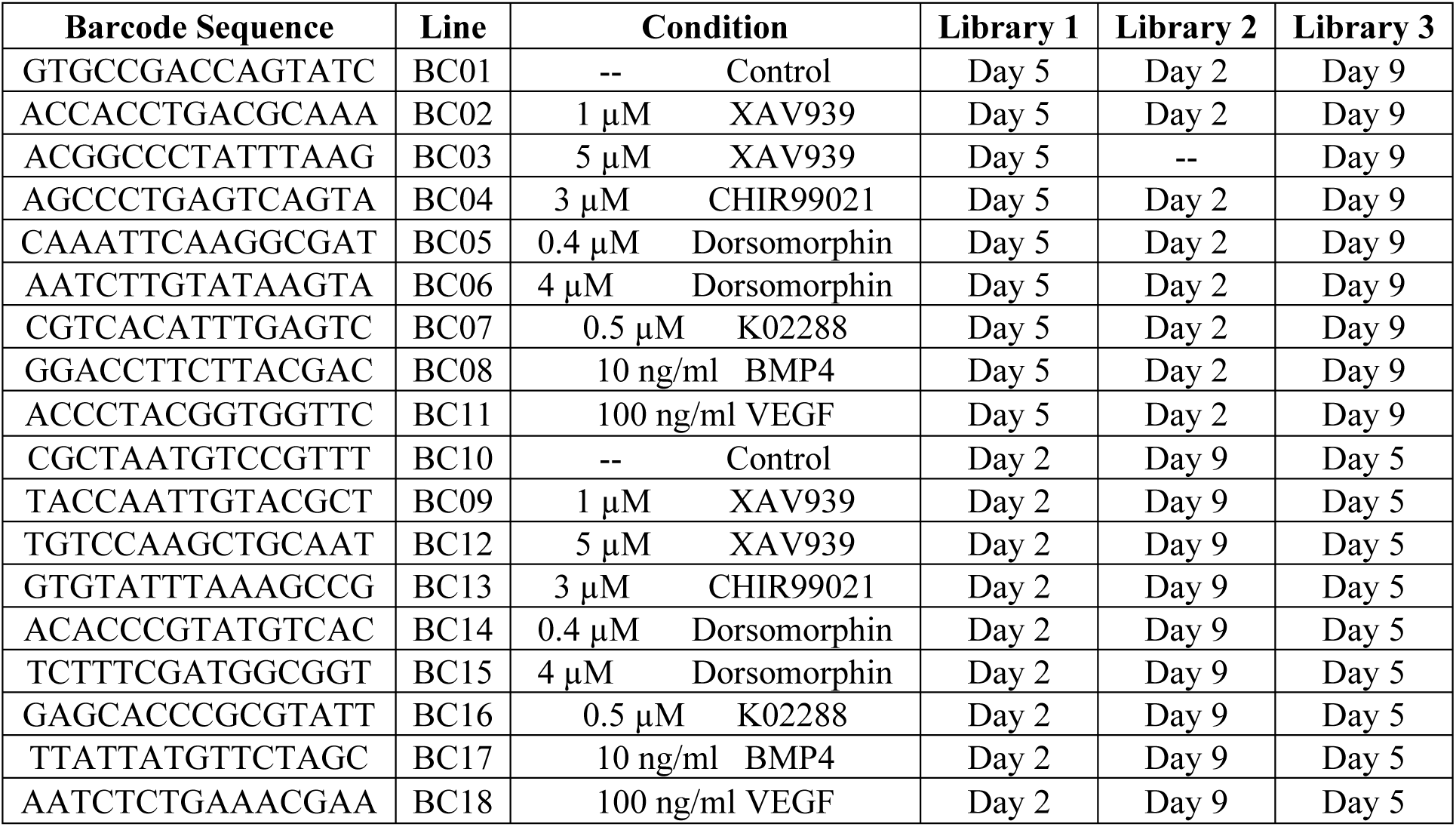
Barcodes and multiplexed experimental design.

We performed a pilot scRNA-seq experiment pooling all 18 barcoding lines as undifferentiated iPSCs to validate homogeneity of their transcriptomic profiles and to confirm that the transcribed barcodes could be used to easily assign a sample of origin for each cell (**Figure 1f**). From a total of 6,711 cells captured, 19.4% had no reads mapped to any genomic barcode, however 95.0% of these cells with no genomic barcode reads also did not pass quality control on the basis of their low total read count, gene count, and high mitochondrial RNA content (**Table S1** and **Figure S1a**). The remainder of cells had reads mapped to between one and seven genomic barcodes, where on average, cells were mapped to two barcodes, with just one read mapped to the lower “expressed” barcode (**Table S1** and **Figure 1g-h**). Dimensionality reduction analysis shows that transcriptomes are sufficiently homogenous to randomly distribute across UMAP space (**Figure 1f**), and cells from each line reveal comparable reads per cell and expression of pluripotency markers *(POU5F1)* (OCT4) and *SOX2* (**Figure 1i** and **Figure S1a**). We additionally used cell hashing antibodies^13^ to label four samples (BC07, BC08, BC12, BC13) to benchmark the accuracy of our method against an established technology. 882 cells were mapped to this subset of genomic barcodes, while 1104 cells had cell hashing (HTO) reads mapped, with an 83.2% agreement between the genomic barcoding and cell hashing sample assignment (**Figure 1j** & **Table S1**). We also observed comparable transcriptomic quality between the two methods (**Figure 1g-j** and **Figure S1a**), providing support for the reliability of using genomic transcribed barcodes for isogenic sample multiplexing in scRNA-seq experiments.

### Generation of a multiplexed atlas of iPSC differentiation *in vitro*

To further evaluate efficacy of these barcoded cell lines for sample multiplexing, we set out to utilise them to capture a dataset that could make full use of the multiplexing potential. Given the diverse signalling and molecular cues shown to inform cell lineage specification during early development, we generated a dataset that could be used to study the role of these molecular determinants during WNT-induced hiPSC differentiation from pluripotency^14,15^ (**Figure 2a**). First, we utilised cell hashing to capture a reference time course of differentiation under control conditions, collecting cells at eight time points; every 24 hours from day 2 to day 9 of differentiation. Next, we treated differentiating cells between day 3 (germ layer differentiation) and day 5 (progenitor cell stage) with perturbations including inhibitors and activators of WNT^16^, BMP^17^, and VEGF^18^ signalling pathways known to be essential regulators of developmental lineage differentiation (**Figure 2b** and **Table 1**). In this signalling perturbation dataset, cells were sampled in duplicates prior to perturbation (day 2), following perturbation (day 5), and at the differentiation endpoint of committed cell types (day 9).

**Figure 2:**
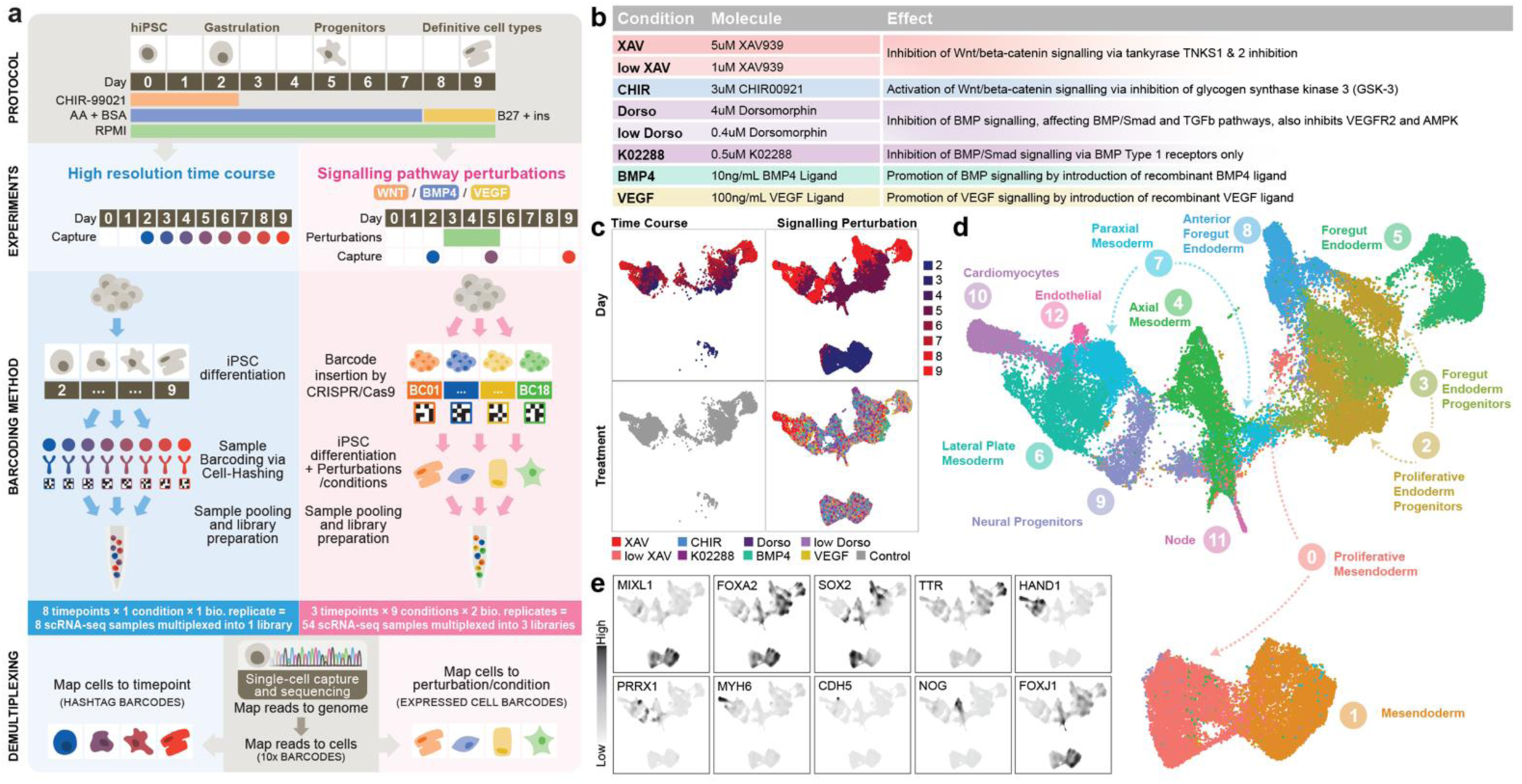
Generating an integrated multilineage atlas of *in vitro* iPSC differentiation. **a)** Schematic of multiplexed study design including 1) a high-resolution time course dataset (blue, left) capturing cells every 24 hours between days 2 and 9 of differentiation and 2) a signalling pathway perturbation dataset (pink, right) generated via perturbation of developmental signalling pathways between days 3 and 5 of differentiation with cells captured on days 2, 5, and 9. In total, 62 cell samples were captured and sequenced using 4 single-cell libraries. Samples were demultiplexed by reference to RNA and sample barcodes. **b)** Signalling perturbation conditions, concentrations, and mechanism of action. **c)** UMAP plots of integrated datasets comprising 62,208 single cells, split by dataset of origin (columns), coloured by time point (top row, 13,682 cells total), and signalling perturbation treatment (bottom row, 48,526 cells total). See also **Table 1**. **d)** UMAP showing the 13 cell type clusters identified using low resolution (0.3) Louvain clustering. Cell type annotations are defined based on expression of marker genes. **e)** Nebulosa^65^ plots showing expression distribution of cell type marker genes. *EOMES* marks mesodermal cells; *SOX2* marks early epiblast, neural progenitors, and foregut progenitors; *FOXA2* marks definitive endoderm; *TTR* marks primitive foregut progenitors*; HAND1* marks precardiac mesoderm; *MYH6* mark cardiomyocytes; *CDH5* marks endothelial cells and *PRRX1* marks limb bud mesoderm. See also **Figure S1f**.

In total, 62 independent cell samples were collected across four scRNA-seq libraries with comparable quality control metrics (**Table S2**). Both the cell hashing library for the time course dataset and genomic barcoding libraries for the signalling perturbation dataset had greater sequencing depth, averaging 600 reads per cell in the former and 190 reads per cell in the latter libraries, where majority of cells had reads mapping to all possible HTO or genomic barcode label (**Table S2** and **Figure S1b**). The cell hashing library showed a greater difference in the percentage of reads mapping to the first and second-highest expressed barcode per cell, however the multiplexing doublet and singlet rates as determined by conventional sample demultiplexing strategies were comparable between the two methods (**Table S2** and **Figure S1b**). Standard quality control and pre-processing of data were performed (**Figure S1c)**, resulting in 13,682 high quality cells from the time course reference and 48,526 cells from the signalling perturbation dataset (**Figure S1d-g**).

The two datasets were integrated using established methods^19^ into an atlas of multilineage hiPSC differentiation (**Figure 2c**). Clustering was performed on the integrated dataset to reveal 13 cell clusters, whose marker gene expression and associated enriched GO terms and KEGG pathways were evaluated to reveal capture of mesendoderm (clusters 0 & 1; *MIXL1*), definitive endoderm (clusters 2 & 3; *FOXA2*) anterior (cluster 8; *SOX2*) and posterior (cluster 5; *TTR*) foregut endoderm progenitors, lateral plate (cluster 6; *HAND1*) and paraxial mesoderm (cluster 7; *PRRX1*), cardiomyocytes (cluster 10; *MYH6*), endothelium (cluster 12; *CDH5*), axial mesoderm (cluster 4; *NOG*), ciliated node cells (cluster 11; *FOXJ1*), and neural progenitors (cluster 9; *SOX2*) (**Figure 2d** and **Figure S2a-c**). Further, expression of pluripotency marker *NANOG* in cluster 0 and enrichment of cell division-related GO terms in cluster 2 distinguish them as proliferative mesendoderm and definitive endoderm respectively (**Figure S2a-c**). Label transfer analysis with reference to mouse^2,20^ and human^4^ embryogenesis scRNA-seq datasets also provided support for these cell type labels (**Figure S2d**). To provide further annotation of the atlas dataset, we used established downstream analysis methods to characterise cell-cell communication between cell types resulting from each treatment condition using CellChat^21^ (**Figure S2e, Table S3**, and **Supplementary File 2**), analysis of enriched gene regulons using pySCENIC^22,23^ (**Figure S2f** and **Table S4**), and URD-predicted^24^ differentiation trajectories relevant to each cluster (**Figure S2g**). Together, this dataset resource presents a deep phenotypic analysis of a time-resolved molecular cell atlas *in vitro* embedded with information of the molecular mechanisms of temporal and signalling control of lineage diversification and cellular heterogeneity during multilineage differentiation of induced pluripotent stem cells.

### Determination of cell subtypes during iPSC differentiation

Following characterisation of the broad cell type groups comprising the dataset, we sought to understand changes in specific cell subtypes from each signalling perturbation. A common approach to this task is to increase the resolution, or granularity, of clustering to delineate smaller cell groups. The problem with this purely algorithm-driven approach is a lack of biological justification for a “correct” clustering resolution or cluster size to select. Given the nuanced signalling cues and small temporal increments in our atlas dataset, we anticipated this cell type ambiguity to be particularly relevant such that increasing clustering resolution would result in many small, uninformative clusters, rather than capturing cell subtypes.

To define such a transcriptional signature marking distinct, more biologically robust cell types, we use a package of tools built from the foundational *TRIAGE* method^25^. In brief, *TRIAGE* is a method that uses consortium-level epigenetic patterns to identify genes predicted to have high cell type-specific regulatory functions. In this way, *TRIAGE* provides an independent biological reference point for defining cell states. This method was adapted for application in single cell analysis pipelines in *TRIAGE-Cluster* and *TRIAGE-ParseR* to help identify cell populations and gene regulatory networks underpinning diverse cell types (**Figure 3a** and **Figure S3a**).

**Figure 3:**
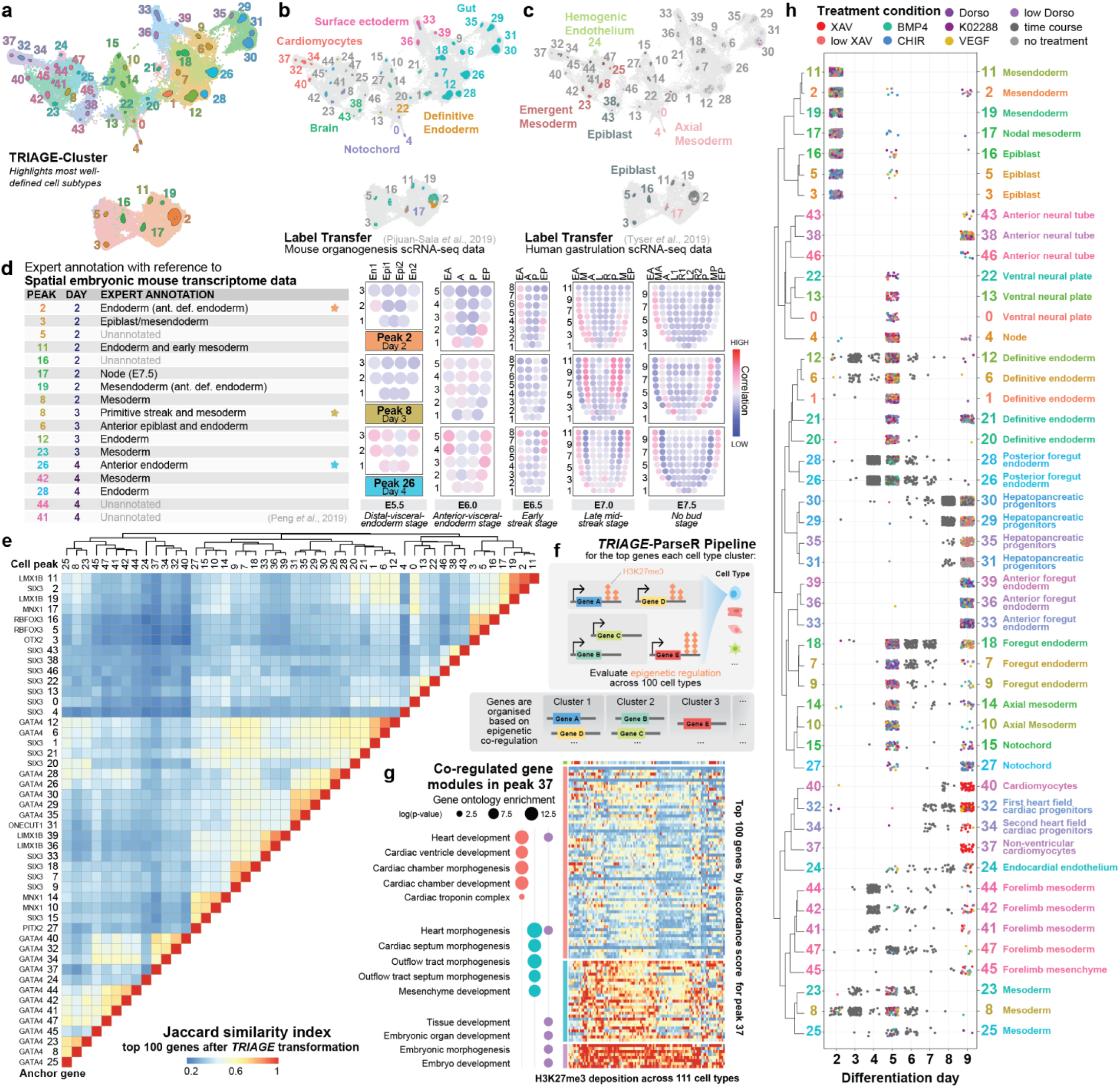
Characterisation and annotation of *in vitro-*derived cell types. **a)** UMAP showing cell type peaks identified from *TRIAGE-Cluster* compared to broad cell type clusters identified in Figure 2d. **b-c)** Annotation of *TRIAGE* cell type peaks by reference to three orthogonal *in vivo* single-cell data sets including (**b**) annotated cell types in single-cell datasets of mouse organogenesis^2^, and (**c**) the gastrulating human embryo^4^. Cells are coloured on a single-cell basis by their assigned label after thresholding (threshold = 0.5). Dark grey cells do not have a label above the threshold. Cell peak number labels are coloured by the label that makes up the greatest proportion of cells in that peak, where grey labels indicate cell type peaks where most cells do not score above the threshold. Light grey cells do not fall within a cell type peak. See also **Figure S3c**. **d)** Annotation of cell peaks by reference to spatial gene expression domains in the developing mouse embryo at five different stages of development between embryonic days 5.5 and 7.5, spanning pre-gastrulation to late primitive streak and amnion formation^27,66^. A Pearson correlation between the expression of selected genes^67^ for each *TRIAGE* peak and each embryonic spatial domain is visualised as corn plots. Annotation of the highly correlated domains for each cell peak and timepoint are summarised in the table (left) and the starred rows’ corresponding corn plots are displayed (right). For each cell peak and timepoint, the five corn plots capture different stages of development in mouse embryos, where each spot has captured the expression profiles of approximately 5-40 cells. Ant. def. endoderm: Anterior definitive endoderm; En1 and En2: divided endoderm; Epi1 and Epi2: divided epiblast; EA: anterior endoderm; A: anterior; P: posterior; EP: posterior endoderm; M: whole mesoderm; L: left lateral; R: right lateral; MA: anterior mesoderm; L1: anterior left lateral; R1: anterior right lateral; L2: posterior left lateral; R2: posterior right lateral; MP: posterior mesoderm. See also **Figure S2d**. **e)** Heatmap and dendrogram showing pairwise Jaccard similarity score between each *TRIAGE-Cluster* cell peak using the top 100 genes by *TRIAGE*-transformation. Anchor genes for each cell peak are listed. **f)** Schematic describing *TRIAGE-ParseR* analysis pipeline^28^ to identify gene clusters from each cell peak to reveal functional and characteristic gene programs of cell subtype. **g)** Example analysis of cardiomyocytes (peak 37) using *TRIAGE-ParseR.* Heatmap shows H3K27me3 deposition across 111 cell and tissue types in the NIH Epigenome Roadmap data across the 100 top ranked genes for cell type peak 37 after *TRIAGE*-transformation. The genes are grouped using the gene co-modulation analysis and GO term enrichment (-log(p value)) of select terms for each of the main gene clusters are shown (left). See **Tables S5-7** for full list of parsed genes and enriched ontologies for all cell type peaks. **h)** Jitter plot showing every cell in the dataset based on the *TRIAGE-Cluster* peak, time point, and signalling condition under-which each cell was derived. Final cell peak annotation is provided (right). Dendrogram (left) reflects pairwise Jaccard similarity of the top 100 genes after *TRIAGE*-transformation^25^ for each cell peak. See also **Figure S3e**.

First, we used *TRIAGE-Cluster* to identify informative cell subtypes within broader cell type clusters, eliminating the need to evaluate tens or hundreds of small, transcriptionally similar clusters (**Figure 3a**, **Figure S3b** and **Table S5**). Based on the clustering output (**Figure 3a**), we took three approaches to evaluate *in vitro-*derived cell peaks by reference to *in vivo* derived developmental cell types. We first used gene set scoring to analyse cells based on positive gene sets defining seven major cell type domains modelled from 921 RNA-seq datasets^26^. We found that among the high-scoring cells, 5 general cell type domains are represented in our data: mesoderm, surface ectoderm, neural crest, germ, and embryonic domains (**Figure S3c**). We also used label transfer analysis to score *in vitro* derived cells based on their transcriptional similarity to single-cell atlases of mouse organogenesis^2^ (**Figure 3b**) and human gastrulation^4^ (**Figure 3c**). These reference datasets provide a preliminary indication of the *in vitro-*derived cell type diversity at the mid-to late timepoints (days 5-9), revealing a range of high-confidence matches to cell types including cardiomyocytes, hemogenic endothelium, and gut progenitor cells (**Figure 3b-c**), aligning with the broad cell type annotations from **Figure 2d**. For the earlier timepoints (days 2-4), we assessed transcriptomic similarity of the cell peaks to embryonic mouse spatial transcriptome data^27^, revealing early cell types corresponding to gastrulation-stage cell types including those in the epiblast, primitive streak, and mesendoderm (**Figure 3d** and **Figure S3d**).

### Evaluation of gene programs underpinning cell subtypes

We next used *TRIAGE* to evaluate cell type-specific gene expression patterns. *TRIAGE* analysis can return a list of the most cell type-defining genes of any given group of cells, akin to differentially expressed or highly variably expressed genes, which can be used as input for common annotation analyses including gene ontology (GO) or KEGG pathway enrichment. (**Table S6**) Unlike differentially expressed genes however, this approach allows each cell group’s expression to be represented without comparison to any other cell cluster in the dataset. We show that the top 100 most highly ranked *TRIAGE* genes for each cell type can be used to inform regulatory relationships between cell types by evaluating shared and unique genes for each cell type peak (**Figure 3e**).

Using these 100 top ranked *TRIAGE* genes for each cell type, we implemented *TRIAGE-ParseR*^28^, which identifies co-regulated molecular processes driving cell type identity (**Figure 3f**). By separating gene lists into related sub-groups, this analysis can enhance enrichment readout from methods such as Gene Ontology (GO) or STRING network analysis and thus clarify cell type annotation (**Tables S6-8**). As an example, *TRIAGE-ParseR* analysis of cell peak 37 reveals three main gene modules with GO enriched terms for cardiac chamber development and morphogenesis of cardiac septa (**Figure 3g**), suggesting its transcriptional profile aligns with the embryonic ventricular septum.

Annotation of cell types was performed by drawing on all the provided results to assign cell type annotations to each *TRIAGE* peak, taking into consideration expression of marker genes, evaluation against *in vivo* reference points, the top *TRIAGE* identity genes, and enriched *TRIAGE-ParseR* gene programs in each cell type. These results provide an integrated view of iPSC *in vitro* multilineage differentiation, to facilitate interrogation of temporal and signalling-dependent cell subtype differentiation (**Figure 3h** and **Figure S3e**).

The atlas reveals cells representing gastrulation stage cell types at day 2 including epiblast, ectoderm, and mesendoderm. Time course data between days 3-5 reveal the emergence of paraxial, cardiac, and forelimb mesoderm together with definitive endoderm and posterior foregut. A sampling of time course and signalling perturbation data on day 5 reveal the first appearance of endocardial endothelium, cells of the anterior foregut, notochord, and neural tube. As cells progress between days 6 and 9 in the time course, we identify the emergence of first and second heart field cardiac lineage cells, and cells of posterior foregut lineages, including pancreas and liver. On day 9, integrated analysis of time course and signalling data show differentiation of additional definitive cell types including ventricular cardiomyocytes, and more developed progenitors of the hepatopancreatic, anterior foregut, and neural lineages. Collectively, these wet and dry lab methods enable the elucidation of mechanisms controlling multilineage differentiation from pluripotency.

### Analysis of signalling pathway-associated cell differentiation to identify genetic regulators of cell fate

We next assessed differentiation lineage biases arising from each signalling perturbation (**Figure 4a** and **Figure S4a**, see also **Figure 2b**). At day 5, cells are broadly distributed across definitive endoderm, foregut endoderm, axial, and general mesoderm. At day 9, however, marked differences emerge. Cells from the control groups show preferential differentiation of posterior foregut-derived hepatopancreatic progenitors, as well as a small proportion of cardiac progenitor cells. This co-differentiation of foregut and cardiac lineages is aligned with embryonic development, where shared signalling cues induce differentiation of the adjacent progenitor populations^29^. Treatments with a lower dosage of XAV-393 (low XAV) and recombinant VEGF produce similar proportions of cell subpopulations to the control at day 9, suggesting that these two treatments are at insufficient levels to affect differentiation at that post-gastrulation mesendoderm stage.

**Figure 4:**
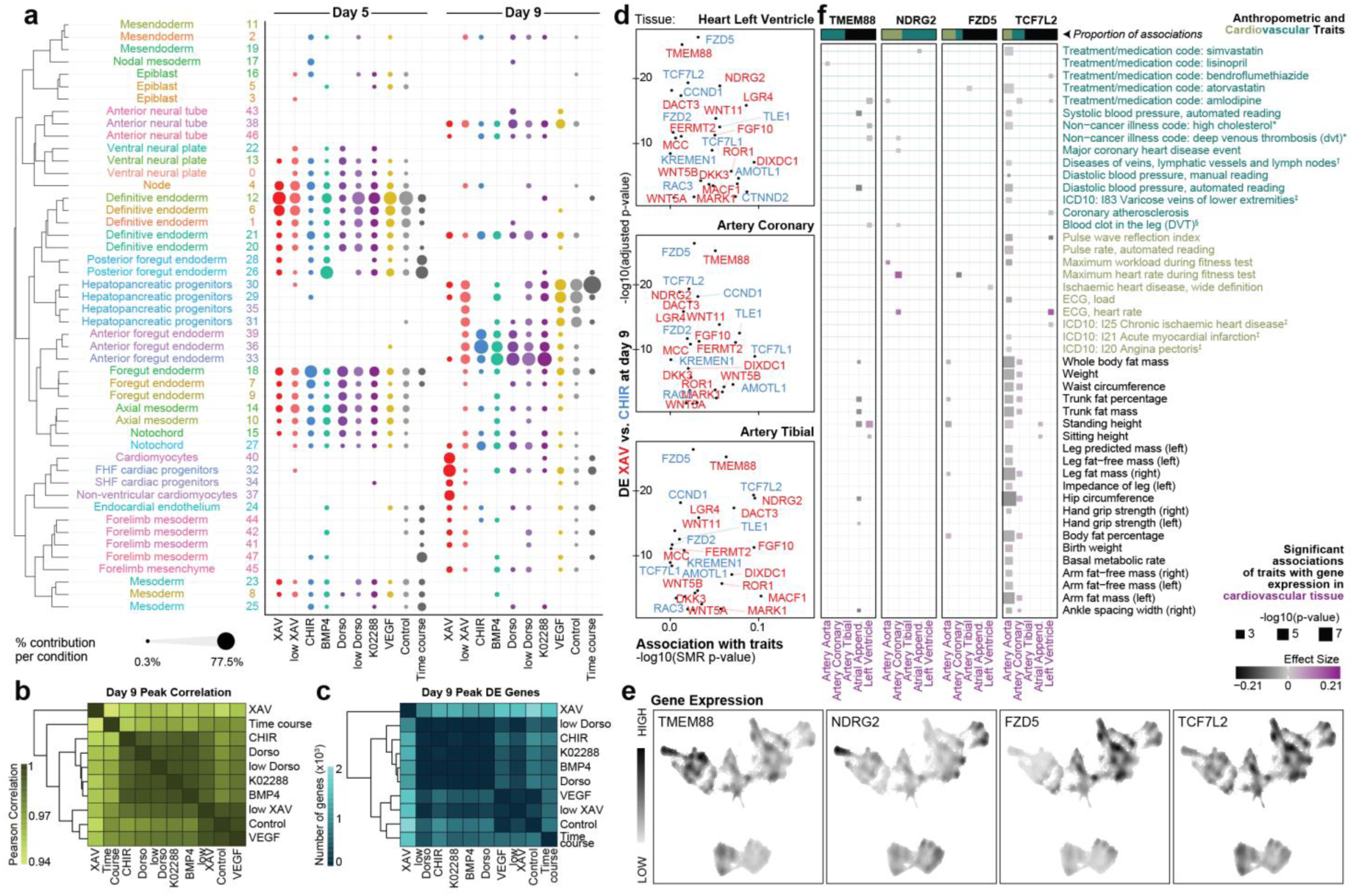
Signalling analysis of cell differentiation. **(a)** Proportion of cells from each treatment condition contributing to each cell type peak at days 5 and 9 of differentiation. Each column represents a treatment condition and sums to 100% across all cells produced by that treatment for the corresponding day of differentiation. FHF: First heart field; SHF: Second heart field. **(b-c)** Heatmaps summarising the pairwise correlation **(b)** and number of differentially expressed genes **(c)** between cells from each treatment condition at day 9. **(d)** WNT-related genes that are differentially expressed between XAV (red) and CHIR (blue) treated *TRIAGE-Cluster* peak cells at day 9 (y-axis) and the significance of the cardiovascular trait most strongly associated with their expression in three cardiovascular tissues, as determined by Summary data-based Mendelian Randomisation (SMR) analysis (x-axis). **(e)** Nebulosa^65^ plots showing expression pattern of four candidate WNT-related genes in the iPSC atlas of mesendoderm differentiation. **(f)** SMR analysis evaluating associations of candidate WNT-related genes’ expression in cardiovascular tissues with cardiovascular and anthropometric traits in the UK Biobank. Some trait and tissue names have been shortened from the following: *non-cancer illness code, self-reported: …; ^†^Diseases of veins, lymphatic vessels and lymph nodes, not elsewhere classified; ^‡^Diagnoses – main ICD10: …; ^§^Blood clot, DVT, bronchitis, emphysema, asthma, rhinitis, eczema, allergy diagnosed by doctor: Blood clot in the leg (DVT); Atrial Append. : Atrial Appendage.

In contrast, differentiation of anterior foregut endoderm was observed in conditions exposed to treatment with the WNT agonist CHIR-99021 (CHIR) as well as all BMP signalling perturbations (**Figure 4a**). Studies have demonstrated an anteriorising role for BMP inhibition in specifying the foregut endoderm, aligning with the observed effect in the three BMP inhibitory treatments (0.4uM and 4uM Dorsomorphin, and K02288 treatments)^30^. For the anticipated posteriorising WNT and BMP4 signalling cues, it is possible that endogenous feedback mechanisms are activated to correct the effects of the CHIR and BMP4 treatments resulting in similar cell type proportions to the BMP inhibitor treatments at day 9. Lastly, **Figure 4a** shows that higher dosage of WNT inhibitor XAV distinctly promoted differentiation of cardiovascular lineage cell types at the expense of the foregut endoderm lineages, in line with established protocols for cardiac-directed differentiation^31^.

To leverage this multilineage atlas to discover genetic factors governing cell differentiation, we first compared differentiated cell types from each signalling perturbation at day 9, evaluating pairwise transcriptional correlation and differential gene expression between treatments (**Figure 4b-c**). These data demonstrate that modulation of the WNT pathway has the most significant impact differentiation trajectory, forming the most distinct cell types by day 9. We therefore used differential gene expression between the two WNT-related treatment conditions affecting on day 9 to identify candidate mediators of WNT-requisite cell fate specification, utilising gene ontology annotations to focus on genes known to influence WNT signalling and its effects (**Figure 4d, y-axis** and **Figure S4b**). This narrowed the list to a set of candidate genes associated with XAV and CHIR modulation of the WNT pathway, primarily mediating cardiac vs. foregut lineage differentiation respectively.

To link these genes to a broader physiological context, we used Summary data-based Mendelian Randomisation (SMR) analysis to evaluate associations between expression of each gene in cardiovascular tissue and effects on genetically-driven anthropometric and cardiovascular traits (**Figure 4d, x-axis**)^32^. We selected the most significantly differentially expressed WNT-related genes in the XAV (*TMEM88* & *NDRG2*) and CHIR (*FZD5* & *TCF7L2*) treatments as candidates for further investigation into specific phenotypes identified from the SMR analysis (**Figure 4d-f**). We found significant associations between eQTL-mediated changes in *TMEM88*, *NDRG2* and *FZD5* expression with myocardial and vascular traits including heart rate and deep venous thrombosis (**Figure 4f**). The bulk of associations for *TCF7L2* stem from variation in its expression in the aorta to affect anthropometric traits including hip circumference and body fat measures, in line with its ties to type 2 diabetes^33^. Given the expression of *FZD5* and *TCF7L2* in the endodermal foregut lineage cell types (**Figure 4e**), we also performed SMR focusing on foregut-related tissue and phenotypes, which showed some enrichment for respiratory phenotypes including asthma (**Figure S4c**).

Of the four candidate genes, *FZD5, TCF7L2,* and *TMEM88* are also significantly upregulated between days 2 and 5 of differentiation, supporting a more direct role during Wnt-driven fate specification at the progenitor stage (**Figure S4d**). *FZD5* and *TCF7L2* have well-established roles in the WNT signalling pathway across foregut, intestinal, and neural development, with the former encoding a receptor in the canonical WNT pathway^34,35^ and the latter a transcription factor effector of WNT signalling^36–38^. *NDRG2* has been shown to inhibit β-catenin target genes in heart, brain and liver development, as well as being relevant in heart failure contexts^39–41^. While prior work has implicated *TMEM88* as a WNT inhibitor relevant in cardiac and pharyngeal pouch development^42–45^, the SMR analysis indicates its expression in cardiac tissue being most relevant to vascular, rather than myocardial phenotypes in human populations (**Figure 4f**). Furthermore, interrogation of organism-wide gene expression databases shows that *TMEM88* expression localises primarily to vascular endothelium in adult organisms (**Figure S4d-e**), leading us to identify *TMEM88* as a candidate with potential for discovery into a broader role in cardiovascular biology.

### Role of *TMEM88* in cardiovascular cell differentiation and physiology

To investigate the role of *TMEM88* on differentiation *in vitro*, we used CRISPRi to generate a doxycycline-inducible *TMEM88* knockdown (KD) hiPSC line (**Figure 5a-b**). We generated a high resolution scRNA-seq time course of mesendoderm differentiation to evaluate the effect of *TMEM88* KD (gain of WNT signalling) compared to control conditions and WNT inhibitory conditions (5µM XAV) (**Figure 5c-d**). We captured 24,841 cells across eight time points between days 2 and 9 of differentiation (**Figure 5d**). UMAP plots show marker genes across the time course dataset and conditions (**Figure 5e**). Using data analysis workflows described in **Figure 2** and **Figure 3**, we identified and annotated 23 cell peaks across the dataset (**Figure 5f-g** and **Figure S5a-e**). Comparing the proportion of cells allocated to each cell type over time for each treatment, we identified notable perturbations in multilineage cell differentiation in *TMEM88* KD cells including persistence of lateral plate and paraxial mesoderm as well as depletion of endothelium and posterior foregut and liver bud progenitor cell types (**Figure 5g**). Genetic validation studies using antibody-based fluorescence activated cell sorting confirmed that, compared to control iPSCs, *TMEM88* KD reduced differentiation of endothelial and cardiac cells *in vitro* (**Figure 5h** and **Figure S5f**), further supporting a role for *TMEM88* in cardiovascular cell identity and differentiation.

**Figure 5:**
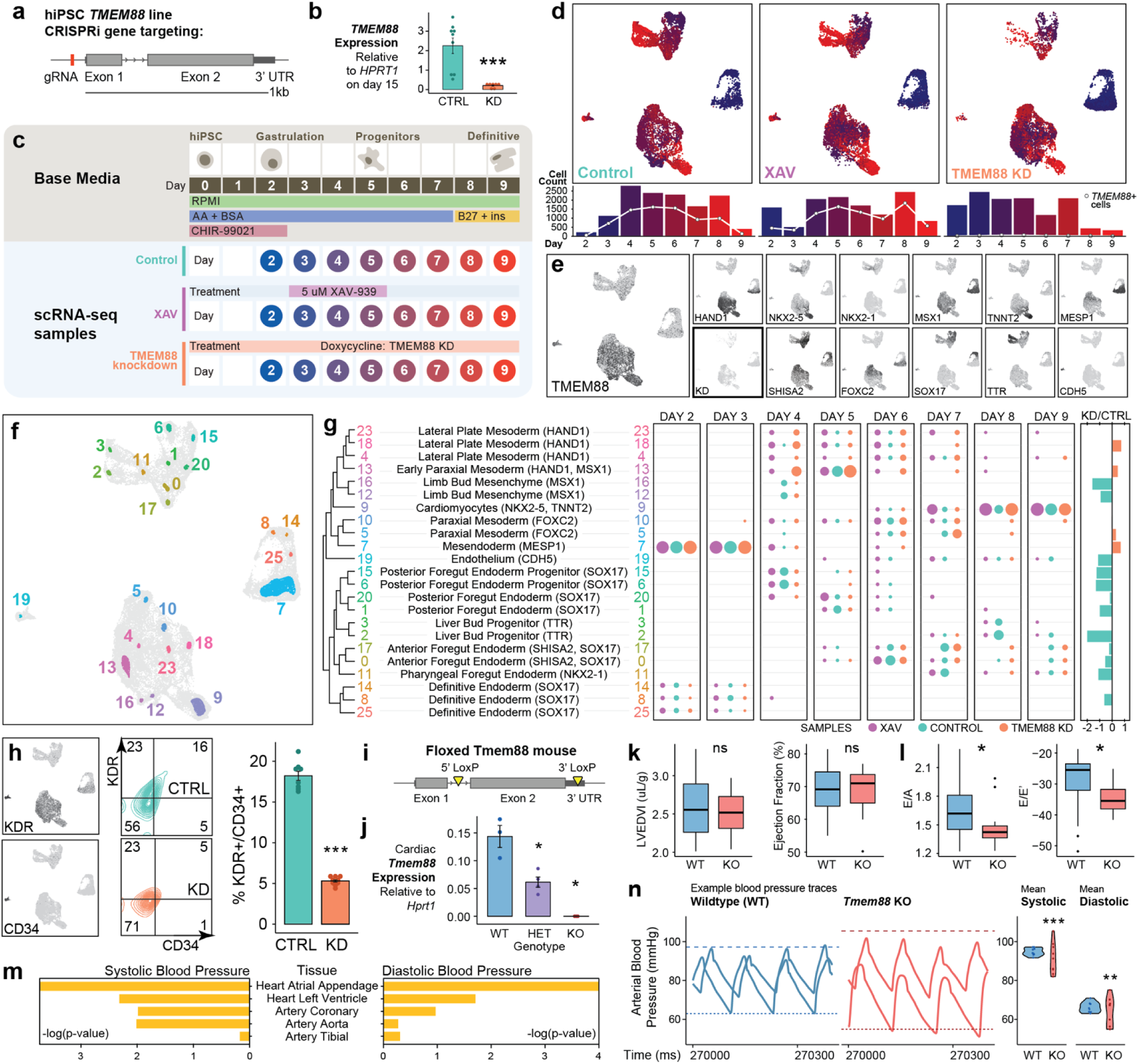
WNT-inhibitor *TMEM88* as a regulator of cardiovascular differentiation and physiology. (a-b) CRISPRi targeting of *TMEM88* to generate doxycycline-inducible *TMEM88* KD hiPS cell line **(a)** and qPCR confirmation of *TMEM88* knockdown efficiency **(b)**, n=7-9, data are represented as mean ± SEM, ***p<0.001 by *t*-test. **(c)** Experimental design of scRNA-seq experiment comparing *TMEM88* KD vs control and XAV-treated cells with cells captured every 24 hrs across day 2-9 of mesendoderm differentiation. **(d)** Distribution of cells at each time point and treatment. Points in the UMAP are coloured by day of collection, and bar plots show number of cells collected at each time point. White points and lines graph overlaid on the bar plot show number of cells expressing *TMEM88* at each time point. **(e)** Expression of marker genes. *TMEM88* (left) in the integrated data set and confirmed loss of *TMEM88* expression in the knockdown sample (KD, bottom row in bold). *HAND1* marks lateral plate mesoderm; *NKX2-5* marks cardiac mesoderm and pharyngeal foregut endoderm; *NKX2-1* marks pharyngeal foregut endoderm; *MSX1* marks paraxial limb bud mesoderm; *TNNT2* marks cardiomyocytes; *MESP1* marks mesendoderm; *SHISA2* marks anterior foregut endoderm; *FOXC2* marks paraxial mesoderm; *SOX17* marks definitive endoderm and its primitive derivatives; *TTR* marks posterior foregut endoderm and *CDH5* marks endothelium. See also **Figure S3c**. **(f)** UMAP showing the 23 cell type peaks identified by *TRIAGE-Cluster* after filtering out those with 20 cells or less. **(g)** Final cell type annotations for cell types identified by *TRIAGE-Cluster* (left) and proportion of cells from each group allocated to each cell type over time (right). Dendrogram (left) reflects pairwise Jaccard similarity of the top 100 genes after *TRIAGE* transformation^25^ for each cell peak. Far right panel quantifies gain or loss of cells in for each cell type peak for *TMEM88* KD vs control. Relevant marker genes are included in brackets next to the cell type annotation. **(h)** Validation of depletion of endothelium from *TMEM88* knockdown *in vitro* by flow cytometry. Expression of endothelial markers KDR and CD34 in the scRNA-seq dataset (left) and bivariate contour plots showing representative flow cytometry analysis (middle) and quantification (right) of control vs *TMEM88* KD for KDR/CD34-labelled cells at day 5 of differentiation, represented as mean ± SEM. n = 8-9 per group. ***p < 0.001 by *t-*test. **(i-j)** Design for CRISPR floxed knockout of *Tmem88* in mice (**i**) and validation of genotype-dependent expression of *Tmem88* in cardiac tissue, represented as mean ± SEM (**j**), n = 3-4 per group. *p < 0.05 by *t-*test. **(k-l)** Echocardiographic analysis of heart function evaluating cardiac parameters by b-mode (**k, left**), m-mode (**k, right**) and pulsed wave doppler (**l**) in 8 week old *Tmem88* KO vs wildtype (WT) mice. Left ventricular end diastolic volume index (LVEDVI) normalised to body weight. n = 13-22, * p < 0.05 by *t-*test. In box plots, centre line shows median, box limits show upper and lower quartiles, whiskers show 1.5x interquartile range, and points show outliers. **(m)** Summary-data based Mendelian Randomisation analysis showing significance of *TMEM88-*associated eQTLs from cardiovascular tissues on systolic and diastolic pressure traits. **(n)** Analysis of blood pressure in the left carotid artery by conductance micromanometry. Raw pressure traces of two example animals per group are provided (left) and mean systolic and diastolic pressures are summarised for *Tmem88* KO versus wildtype (WT) mice. Dashed line indicates the highest mean systolic blood pressure and dotted line the lowest mean diastolic blood pressure per group. n = 9, ** p<0.01 and *** p<0.001 by Bartlett test for homogeneity of variances after Shapiro normality test.

To evaluate the role of *TMEM88* in cardiovascular development *in vivo,* we generated a *Tmem88* conditional knockout mouse model by crossing a floxed *Tmem88* mouse (**Figure 5i**) and a mouse with ubiquitous Cre recombinase expression driven from a CMV promoter. Quantitative RT-PCR analysis of heart tissue showed genotype-dependent loss of *Tmem88* gene expression (**Figure 5j**). All genotypes were born in Mendelian ratios and displayed no observable gross developmental phenotypes (**Figure S5g**). However, micro-computed tomography scans of E15.5 embryos showed significant changes to organ volumes and structures including the vertebrae, pelvic girdle, and ductus arteriosus in *Tmem88* KO embryos (**Figure S5h**).

Focusing on cardiovascular physiology, analysis by two-dimensional b-mode and one-dimensional m-mode echocardiography showed no impacts on cardiac structure and function in adult mice (**Figure 5k** and **Figure S5i**), suggesting a minor role for *Tmem88* in cardiac morphogenesis and function. Doppler analysis of blood flow across the mitral valve showed a significant reduction in the E/A and E/E’ in *Tmem88* KO mice, suggesting mild diastolic dysfunction (**Figure 5l**). Lastly, based on SMR results indicating significant associations between *TMEM88* and blood pressure phenotypes across diverse tissues (**Figure 5m** and **Table S9**), we evaluated systemic arterial blood pressure in WT and KO littermates using catheter-based conductance micromanometry analysis of pressure traces from the right carotid artery (**Figure 5n**). These data show that, compared to WT controls, *Tmem88* KO mice had significantly increased variability in systolic and diastolic blood pressure, a phenotype associated with diverse clinical pathologies including hypertension and end-organ damage^46^. To our knowledge, this is the first data implicating *TMEM88* in developmental regulation of blood pressure, providing compelling genetic evidence of its tissue specificity and functional significance that justify further work to understand the biological mechanism of action. Collectively, these data demonstrate the power of coupling large-scale multilineage iPSC differentiation *in vitro* with population-level human statistical genetic analysis of complex traits as a basis to discover genetic regulators of developmental physiology.

## DISCUSSION

This study generated barcoded hPSCs to facilitate multiplexed scRNA-seq analysis of cell differentiation, demonstrated by generation of a single-cell transcriptomic atlas chronicling differentiation of pluripotent stem cells *in vitro*. We further implement a two-part computational approach to cell clustering, including mapping to *in vivo* cell types and classification of well-defined cell subtypes across diverse lineages to provide, at single-cell level, the molecular attributes of multilineage cell differentiation. We couple these data with human population-level complex trait data to link candidate genetic regulators of development with insights into putative physiological functions in adult mammals. Using genetic models, we validate a role for *TMEM88* in governing cardiovascular development, with a primary impact on regulating mammalian blood pressure.

This study provides diverse new tools and harnesses data to maximise knowledge-gain into mechanisms of development and disease. First, we develop novel cell barcoding strategies. Unlike existing scRNA-seq sample multiplexing strategies^13,47,48^, endogenous cellular barcodes broaden the experimental capability for *in vitro* and *in vivo* studies of hPSC differentiation^49^. Barcoding isogenic lines via gene targeting to the *AAVS1* locus allows for constitutively expressed barcodes in all cell types for direct comparison of perturbations, reducing variability related to reprogramming and genotype differences between hPSC^50,51^ samples and simplifying sample preparation. The CRISPR-Cas9 integration of stable genomic barcodes in this study notably differs from barcoding methods that utilise viral barcode delivery into cells for isogenic multiplexing and lineage tracing applications^49,52^. Lentiviral barcode delivery is highly flexible and allows for successive rounds of barcode incorporation over the course of an experiment to allow direct capture of ancestor-descendant relationships during differentiation and reprogramming^53^. As the genomic barcodes are introduced prior to the start of an experiment, they are limited to lower throughput clonal tracking experiments, or those where separately-prepared and barcoded progenitor cells are pooled for co-culture. This method however has the benefit of a defined barcode integration location, minimising potential for off-target effects and more consistent expression across cells and cell types. This proves beneficial in contexts involving high cell type heterogeneity or significant changes in cell states, where lentiviral barcodes are more susceptible to cell type-specific silencing. Overall, as pre-experimental sample multiplexing methods, both barcode delivery methods have complementary use-cases and allow scalable, complex experimental designs for high throughput protocol development, organoid modelling, and *in vivo* cell transplantation studies.

Cell type annotation remains one of the most challenging issues in single-cell studies due to the need to relate cell heterogeneity to known biology. In this study, we implement a computational analysis pipeline outlined in a package of tools (*TRIAGE*^25^, *TRIAGE-Cluster*, and *TRIAGE-ParseR*^28^) to define evolving cell types *in vitro* by referencing identity-defining genetic features. By restricting the analysis to biologically grounded cell types, we eliminated noise arising from technical variability in scRNA-seq capture and hPSC culture which complicate data interpretation. We demonstrated that this method facilitates more precise predictions of cell identity relationships. Furthermore, while machine learning approaches for cell type annotation are increasingly popular due to their automatic, unsupervised classification of cell types^54^, they rely heavily on prior knowledge of cell type annotations. This is problematic particularly among intermediate cell types such as those derived at early stages of *in vitro* differentiation from pluripotency. This study demonstrates that *TRIAGE* provides an unsupervised pipeline to evaluate cell types in single-cell data atlases spanning developmental and mature cell types.

Transcriptional analysis of *in vitro* multilineage cell diversification provides insights into genetic regulation of cell decisions but has limited capabilities for interpreting tissue-level physiological attributes. We demonstrate the value of complementing *in vitro* atlas data with complex trait genetic data from adult human populations to inform physiological studies of cardiovascular development and disease. In particular we highlight the potential of this approach to facilitate discovery of new mechanisms and roles for minimally-characterised genes such as *TMEM88*. While we identify *TMEM88* as a WNT antagonist that regulates cell fate specification, prior studies have shown that WNT inhibition treatment only partially rescues the effects of its loss-of-function in cardiac development, suggesting possible function through other pathways unrelated to WNT signalling in development. Further, given the mismatch of phenotypes between *in vitro* and *in vivo TMEM88* loss-of-function likely arising from compensatory mechanisms masking more subtle or complex phenotypes *in vivo*^44,55^, reference to independent statistical genetics data provided a novel hypothesis for *TMEM88* in an adult context. Our validation of this role *in vivo* reinforces the relevance of studying mechanisms that regulate lineage differentiation in broader developmental and disease contexts^56^. This study and approach therefore provide evidence for a novel drug target to address the ongoing health burden of cardiovascular disease.

Blood pressure variability is a significant risk factor in organ damage and a major determinant for diseases including atherosclerosis, chronic kidney disease, and dementia^46,57,58^. Though it is more common among the 33% of adults experiencing hypertension globally^59^, blood pressure variability is independently associated with cardiocerebrovascular events including stroke and myocardial infarction, the two leading causes of death worldwide^60,61^. Analysis of one of the world’s largest population-based assessments of middle-aged adults with measured blood pressure found that most treated hypertensives remain uncontrolled^62^, and only a subset of antihypertensive drugs have demonstrated capability to target both high and variable blood pressure^63^. As there are also no specific treatments for blood pressure variability^64^, identification of novel gene targets governing blood pressure, such as *TMEM88*, provides new opportunities to address this persistent public health concern.

## LEAD CONTACT

Further information and requests for resources and reagents should be directed to and will be fulfilled by the Lead Contact, Nathan J. Palpant

## ACKNOWLEDGEMENTS

We thank Nick Valmas for illustrations; the Australian Genome Research Facility (AGRF) for Sanger sequencing; the Flow Cytometry Facility at the Queensland Brain Institute for facilitating cell sorting; the University of Queensland sequencing facility at the Institute for Molecular Bioscience for performing single-cell capture and library preparation; and the Garvan Sequencing Platform for sequencing. The mutant floxed *Tmem88* mice were produced via CRISPR genome editing by the Monash Genome Modification Platform, Monash University as a node of Phenomics Australia (formerly Australian Phenomics Network). Phenomics Australia is supported by the Australian Government Department of Education through the National Collaborative Research Infrastructure Strategy, the Super Science Initiative, and the Collaborate Research Infrastructure Scheme. We acknowledge the Victor Chang Research Institute Innovation Centre, funded by the New South Wales Government.

Funding support was provided from the National Health and Medical Research Council of Australia (Grants 1143163 (N.J.P., J.E.P., and S.W.L.), 1110751 (P.P.L.T.), 1173469 (P.Y.), GNT2008928 (Q.N.), APP1107599 (J.E.P.), 2007896 (S.L.D)), the Medical Research Future Fund (APP2016033, N.J.P. and M.B.), the Australian Research Council (190102793 (P.P.L.T.)), the National Heart Foundation of Australia (Grants 101889 and 106721, N.J.P.), and AIR@InnoHK administered by Innovation and technology Commission (J.W.K.H).

## AUTHOR CONTRIBUTIONS

To generate the barcoded iPS cell lines, S.L. designed the barcodes; S.A. and D.X. designed the plasmids. S.A. cloned the plasmids; T.W. and H.C. carried out the cell culture; D.X. designed the CRISPR targeting strategy. T.W. and D.X. performed the CRISPR gene editing. M.T. and D.X. performed the genotyping and Sanger sequencing. For the pilot barcoding scRNA-seq, T.W. and H.C. carried out the cell culture and collection; S.L. and S.A. processed and sequenced the samples; S.L. and S.S. performed the computational sample demultiplexing and quality control analysis. J.E.P. supervised the barcoding design and assisted with sequencing of barcoded libraries and facilitated data interpretation.

For the time course and *TMEM88* KD scRNA-seq datasets, H.C. generated the CRISPRi hiPSC line; S.S. and X.C. carried out cell culture, collection, and sample preparation; S.S. performed quality control and preliminary computational analysis. For the signalling perturbation scRNA-seq dataset, T.W. and H.C. carried out cell culture, collection, and sample preparation; S.S. and T.W. completed quality control and preliminary computational analysis. For analysis of the combined dataset, S.S. integrated the data and led analysis; E.S., Q.Z., and Y.S. performed additional characterisation analysis of broad cell clusters; Y.S. and S.S. conceptualised and implemented the *TRIAGE*-*Cluster* approach; Y.S. developed and benchmarked the *TRIAGE-Cluster* method with supervision from W.S.; D.K., P.Y., and P.P.L.T. performed annotation of cell types with reference to spatial gene expression patterns in the developing mouse embryo; D.P. constructed the online dataset visualisation dashboard and website; Z.S. implemented the visualisation of gene expression to the platform; Q.N. supervised creation of the website; J.W.K.H supervised implementation of the gene expression visualisation; W.S. conceptualised and implemented the *TRIAGE*-*ParseR* analysis pipeline; M.B. supervised development of the *TRIAGE*-*ParseR* software.

For analysis of the role of *TMEM88*, D.M. performed statistical genetics analysis of *TMEM88* on human complex trait data; S.N. provided preliminary analysis of GWAS data linking variants in *TMEM88* with vascular physiology; S.K. performed analysis of *TMEM88* KD cardiac and endothelial iPS-derived cell types; E.M.M.A.M. performed micro-computed tomography and data quality control; G.D. performed LAMA analysis micro-computed tomography datasets and interpreted the results; S.L.D. supervised micro-computed tomography and LAMA analysis of the *Tmem88* embryos; S.S. performed echocardiographic imaging and associated analysis with supervision from M.A.R. and J.O.; J.O. performed conductance micromanometry analysis for analysis of blood pressure.

N.J.P. conceptualised the study, supervised the project, and raised funding. S.S. and N.J.P. wrote the manuscript.

## STAR METHODS

### hiPSC culture and standard monolayer mesendoderm differentiation

All human pluripotent stem cell studies were carried out in accordance with consent from the University of Queensland’s Institutional Human Research Ethics approval (HREC#: 2015001434). Undifferentiated hiPSCs were cultured on Vitronectin XF (STEMCELL Technologies #07180) coated plates in mTeSR1 media (STEMCELL Technologies #05850) with supplement at 37°C with 5% CO_2_. On the day prior to differentiation (day −1), cells were dissociated using 0.5mM EDTA solution and seeded onto separate coated plates in pluripotency media with ROCK Inhibitor (STEMCELL Technologies #72308) and cultured overnight. Once forming an ∼80% confluent monolayer, differentiation was induced (day 0) by changing the culture media to RPMI (ThermoFisher, #11845119) containing 3µM CHIR99021 (STEMCELL Technologies, #72054), 500mg/mL BSA (Sigma #A9418), and 213mg/mL ascorbic acid (Sigma #A8960). On days 3 and 5, the media was replaced with the same media cocktail excluding the CHIR99021 supplemental cytokine. On day 7 and every subsequent other day, the cultures were fed with RPMI containing 1xB27 (ThermoFisher #17504001) supplement plus insulin.

### Barcode cassette design

The barcodes (**Table 1**) were introduced into the cells as a part of a barcode cassette, which also incorporated the reverse complement of a partial Chromium Read2 adaptor sequence (truncated slightly to allow oligo length of <= 60bp). This Read2 sequence is used as a PCR handle to generate barcode-containing amplicons compatible with Chromium scRNA library preparation. Additionally, restriction enzyme recognition sequences were added or regenerated to enable easy transfer of the cassette between different vectors. The exact structure of the cassette is as follows: EcoRV site 3’ 3bp – MluI site – partial 10x Read2 adaptor reverse complement – 15bp barcode – MluI complementary sequence. Barcoding cassettes were ordered as complementary single-stranded oligos, which could be annealed and ligated into a digested plasmid backbone.

### Vector design

AAVS1-CAG-hrGFP (Addgene #52344) was used as the plasmid backbone. It contains hrGFP under the control of the CAG promoter, and AAVS1 homology arms to allow precise integration by homologous recombination of the donor sequence into the *AAVS1* locus of the genome when paired with the CRISPR/Cas9 system using a well-described guide RNA^68^. The barcode cassette was introduced between the EcoRV and MluI sites of the plasmid between hrGFP and the polyadenylation site, to enable expression as part of the hrGFP transcriptional unit (**Supplementary File 1**).

### Generation of barcoding plasmids

AAVS1-CAG-hrGFP (Addgene #52344) was digested by incubation with EcoRV-HF (New England BioLabs; NEB #R3195S) followed by addition of MluI-HF (NEB #R3198S) and further incubation. Successful digestion was confirmed by running a small amount on an agarose gel, and remainder was purified using QIAquick PCR Purification Kit (QIAGEN #28104). Top and bottom strands of barcode oligos were annealed by mixing 1uL each of 100 µM oligo in 1x T4 DNA ligase buffer in a volume of 10 µL, heating to 94°C for 2 min, then allowing to cool to 25°C at a rate of 1°C/s. Annealed oligos were further diluted 1 in 10 with nuclease-free water. Ligation of barcode cassettes to vector was performed by combining 100ng digested plasmid, 4 µL diluted annealed oligos, and 1 µL T4 DNA ligase (NEB #M0202S; 400U/µL) in 10µL total volume with 1x T4 DNA ligase buffer. Reaction was incubated at 16°C for 16 hours. Separate reactions were performed for each barcode oligo. 3µL of ligation reactions were added to 20 µL Stellar competent cells (Takara #636763) and heat shock transformation performed at 42°C for 1 min in 1.5mL tubes. 350 µL SOC media was added for recovery, with shaking at 37°C for 1 hour. 100 µL was spread onto selective ampicillin-containing agar plates, which were incubated overnight at 37°C. 3 colonies were picked from each plate for screening with colony PCR for expected insert in 10 µL reactions using MangoTaq (Bioline #BIO-21083). Colonies showing successful amplification of insert were grown overnight in 5 mL LB broth containing ampicillin, and plasmid purified using QIAGEN Plasmid Miniprep kit. 100% sequence identity was confirmed across the barcode insert by Sanger sequencing using a universal sequencing primer (5’-ttttggcagagggaaaaaga-3’), performed by the Australian Genome Research Facility (AGRF). 50 mL cultures were grown from glycerol stocks of plasmids with confirmed barcode sequence insert, and plasmid was purified using Nucleobond Xtra Midi Kit (Macherey-Nagel #740410.50) to give endotoxin-free, high concentration stock for transfection.

### Stable generation of WTC BC01-BC18 barcoded iPS cell lines

For gene editing, WTC WT-11 hiPSCs (Gladstone Institute of Cardiovascular Research, UCSF; Karyotype: 46, XY; RRID: CVCL_Y803, generated as previously described^69,70^) cells were grown to about 50-80% confluency, dissociated using 1XTrypLE and 100-200K cells were used for each 10 µl reaction of the Neon Transfection System. The transfection mixture included 0.5 µg of barcoding plasmid DNA, 20 pmol AAVS1-taregting sgRNA (protospacer sequence: GGGGCCACTAGGGACAGGAT, chemically synthesised by Agilent technology) and 20 pmol spCas9 protein (IDT). After electroporation with 1 pulse of 1300 V for 30 ms, cells were seeded in mTeSR1 with ROCK Inhibitor (Y-27632; STEMCELL Technologies #72308) and CloneR (STEMCELL Technologies #05888). Selection was performed with 1 µg/ml puromycin, and purified cell lines were frozen down in CryoStor CS10 Cell Freezing Medium (STEMCELL Technologies #07930) and stored in liquid nitrogen.

### Quality control of cell lines

All cell lines underwent quality testing for correct genetic insertion, selection efficiency, pluripotency, chromosomal abnormalities, and mycoplasma contamination. Genomic DNA from all cell lines was extracted using QuickExtract DNA Extraction Solution (Epicentre #QE09050). Correct targeting of donor construct at the *AAVS1* locus was confirmed by junction PCR using the following primer pair: AAVS1 F1: 5’-ggttcggcttctggcgtgtgacc-3’, AAVS1 R1: 5’-tcaagagtcacccagagacagtgac-3’. The purified PCR product was then Sanger sequenced using a universal sequencing primer (5’-cccatatgtccttccgagtg-3’) to validate correct barcode insertion in each cell line. Flow cytometry was performed on live cells for endogenous GFP expression and after labelling for the pluripotency marker SSEA3 (BD Biosciences #562706) and a corresponding isotype control. Karyotyping was carried out as a professional service by Sullivan Nicolaides Pathology. iPSCs were grown in a 25 cm^2^ flask to about 70-80% confluency and sent for analysis, 15 cells were examined per culture.

### Single-cell RNA-sequencing of undifferentiated barcoded iPSCs

All barcoded iPS cell lines were cultured in parallel as described above, dissociated using 0.5mM EDTA and 600K cells from each cell line were combined. Prior to this, four cell lines were additionally labelling with different TotalSeq-A cell hashing antibodies^13^ according to the manufacturer’s protocol. The combined sample was transferred into 2% BSA (Sigma #A9418) in PBS, stained with Propidium Iodide and 500K viable cells were sorted using a BD Influx™ Cell Sorter (BD Biosciences) with FACSDiva software (BD Biosciences).

Single-cell RNA-seq libraries were generated using the 10X Genomics Chromium Single Cell 3’ v2 protocol, with minor modifications to the workflow, outlined by Stoeckius and Smibert to capture the fraction of droplets containing the Hashtag Oligonucleotide (HTO)-derived cDNA (<180bp). HTO additive primers and Illumina TruSeq DNA D7xx_s primer (containing i7 index) were ordered from IDT, and used according to the cell hashing protocol. Hashtag libraries were quantified using the Agilent Bioanalyzer. Sequencing was performed using the Illumina Nextseq instrument, with gene expression and HTO libraries pooled on a single flow cell using a ratio of 90:10. Raw sequencing data were processed using the the 10X Genomics Cell Ranger pipeline to derive gene expression count matrices.

HTO-tagged cells were identified and extracted from the fastq files using the CITE-seq-Count^71^ program with default parameters, to generate a count matrix of cells and their respective HTO expression values. This allowed the pooled hashing cells to be independently deconvoluted. From the 3’ gene expression data, barcoded cells were identified by the expression of a barcode from the whitelist, which was included in the transcriptome reference with unique identifiers (e.g., ‘BC01’).

Transcriptome-based quality control filtering was performed, keeping cells with library sizes between 7500 and 60,000 reads, between 2000 and 8000 distinct features, mitochondrial reads making up no greater than 20% of all reads, and ribosomal reads mapped to between 22 and 45% of total reads. The *scds* R package^72^ was also used to remove doublets predicted based on transcriptomic features. For this pilot study, a simplistic demultiplexing method for both the genomic barcodes and the cell hashing multiplexing was used where the highest barcode was taken to be the sample identity for each cell, cells with no counts mapped to any barcode were assigned as ‘Negative’ and cells with two or more barcodes expressing the same maximum count were determined to be ‘Doublets’ or ‘Multiplets’. The agreement between genomic and hashing barcode assignments used is the sum of the double positives and double negatives divided by the total number of cells. Standard normalisation, scaling, and dimensionality reduction of this dataset was done using the *Seurat* R package to generate the plots.

### Cell hashed high resolution time course of mesendoderm-directed differentiation

The cell line used to generate the time course and *TMEM88* scRNA-seq datasets were WTC CRISPRi TMEM88-g2.3 GCaMP hiPSCs (Karyotype: 46, XY; RRID: CVCL_VM38; generously provided by M. Mandegar and B. Conklin, Gladstone Institute, UCSF), generated as previously described^73^. In brief, guide RNA targeting the transcriptional start site of *TMEM88* in the human genome was cloned into the pQM-u6g-CNKD doxycycline-inducible construct as previously described^73^ (Forward oligo: 5’-TTG GAG AGC CGC ATT CCA GGA TTA-3’; Reverse oligo: 5’-AAA CTT ATC CTG GAA TGC GGC TCT-3’). The construct was then transfected into WTC CRISPRi GCaMP hiPSCs using the GeneJuice (Novagen), followed by selection using rounds of replating with 10ug/mL blasticidin treatment. Populations were tested for knockdown efficiency of *TMEM88* (Forward primer 5’-CCT ACT GGT CAC CGG ATT CCT-3’, Reverse primer 5’-GAC GCC GAT AAA GGG CTC G-3’), normalised to housekeeping gene *HPRT1* by qPCR (see also **Quantitative RT-PCR**) following continuous 1ug/mL doxycycline treatment from day 0 of differentiation.

For the time course experiment, cells were not exposed to doxycycline and treated as transcriptionally wildtype. Mesendoderm-directed differentiation as described above was induced in separate plates for each collection timepoint. Cells were dissociated using 0.5mM EDTA in 2.5% Trypsin (ThermoFisher, #15400054) and neutralised with 50% foetal bovine serum (GE Healthcare Life Sciences, #SH30084.03) in DMEM/F12 media (Sigma #11320033). 1 x 106 cells from each sample were labelled with a different TotalSeq-A cell hashing antibody (BioLegend antibodies A2051-8) as per the recommended protocol ^13^ and sorted for viability on a BD Influx Cell Sorter (BD Biosciences) using propidium iodide. 5 x 105 live cells per time point were collected and pooled for Chromium Single Cell 3’ V3 (10x Genomics) reactions following the manufacturer’s protocol, targeting 2×10^4^ cells. Gene expression libraries were sequenced on an Illumina NovaSeq 6000 and the cell hashing libraries were sequenced on an Illumina NextSeq 550Dx.

### Multiplexing of signalling perturbations during mesendoderm differentiation

18 barcoded iPS cell lines (WTC BC01-BC18) were cultured in parallel and multiple staggered set-ups of mesendoderm-directed differentiation. Cell lines were divided into 2 batches (BC01-BC09 and BC10-BC18) to allow for the capture of biological duplicates for all 9 conditions. Cells were collected for scRNA-seq on day 2 as a reference prior to any perturbations, as well as on days 5 and 9 of differentiation. In total, 3 sequencing libraries were generated, with each library consisting of all 18 cell lines. A combination of 2 different timepoints from the two batches was pooled in each library to allow for easy detection and removal of potential batch effects during downstream analysis (**Table 1**).

### Signalling pathway perturbation experiment single-cell library preparation and sequencing

Sample pools were assessed for quality using a hemocytometer with Trypan Blue exclusion. Cell viability ranged from 82-88%; cell concentration was between 1.3E+06 and 2.1.9E+06 cells/mL. Chromium Single Cell 3’ v3 (10x Genomics) reactions were performed for each sample according to the manufacturer’s protocol, targeting 20,000 cells per reaction. 11 cycles of cDNA amplification were performed in a C1000 Touch thermocycler with Deep Well Reaction Module (Bio-Rad). After clean-up of full-length amplified cDNA, 10µL was used for the construction of the gene expression library according to the manufacturer’s protocol, with 11 indexing PCR cycles. Additionally, 5µL of full-length amplified cDNA was used to generate a barcoding library for each sample pool. Briefly, a first round of PCR was performed to specifically amplify cDNA regions containing the barcode cassette and append partial P5 and P7 sequencing adaptors. Each reaction contained 1x KAPA HiFi HotStart ReadyMix and 300nM each barcode_amp_F (5’-ACT GGA GTT CAG ACG TGT GCT CTT CCG ATC T-3’) and barcode_amp_R (5’-CTA CAC GAC GCT CTT CCG ATC T-3’) primers in a final volume of 50µL. A 2-step PCR protocol was performed with annealing/extension at 71°C for 30s, for six cycles. After a 1.2X SPRI clean-up to remove primers, a second round of PCR was performed with the entire volume of purified product from PCR1. Each reaction contained 1x KAPA HiFi HotStart ReadyMix, 500nM SI-PCR primer (5’-AAT GAT ACG GCG ACC ACC GAG ATC TAC ACT CTT TCC CTA CAC GAC GCT C-3’), and 5µL of a unique i7 indexed R primer from Chromium i7 Multiplex Kit (10x Genomics), in a total volume of 50µL. PCR was performed as for the SI-PCR protocol in the Chromium gene expression library construction workflow. Eight indexing PCR cycles were performed, for a total of 14 cycles over two rounds of PCR. Final barcoding libraries were purified using 1X SPRI beads, and fragment size and library concentration verified along with gene expression libraries using a BioAnalyzer DNA High Sensitivity Kit (Agilent). Final gene expression libraries were 62-82nM, with average size 457-494bp. Barcoding libraries were 30-38nM, with average size 358-362bp.

A single pool was prepared from the three gene expression and three barcoding libraries for sequencing. The samples were pooled equimolar within each library type and combined so that the gene expression libraries together made up 90% of the pool, and the barcoding libraries 10%. Samples were sequenced on an Illumina NovaSeq 6000 (NovaSeq control software v1.6.0/Real Time Analysis v3.4.4) using a NovaSeq 6000 S4 Kit V1 (200 cycles) (Illumina, 20027466) in standalone mode as follows: 28 bp (Read 1), 8 bp (I7 Index), and 91 bp (Read 2). Gene expression count matrices were derived using the standard 10X cell ranger pipeline.

### Demultiplexing and quality control

Barcoding reads used to assign sample barcodes to each cell were from the separately amplified and sequenced barcode libraries, and cell barcodes without both transcriptome and sample barcoding reads were removed. For each barcode sequencing library, the ‘HTODemux’ function in the *Seurat* R package^74^ with the default 0.99 quantile cutoff was used to determine the dominant sample barcode for each cell and annotate negative and doublet cells based on their sample barcode reads alone. Three transcriptome-based doublet detection methods in the *scds* R package were used to further assign doublet annotations to each cell, and cells labelled as doublets by at least three methods were removed. Transcriptome-based cell filtering as part of the *Seurat* pipeline removed cells with fewer than 2000 and greater than 7500 detected genes; fewer than 5000 and greater than 50,000 total read counts; or mitochondrial reads accounting for greater than 25% of total reads. Following filtering, sample barcodes were assigned to the remaining cells based on the barcode with the highest expression in each cell.

### Visualisation of individual scRNA-seq datasets

Normalisation and UMAP dimensionality reduction of the data were done following the standard pipeline in the *Seurat* pipeline. To visualise the expression of marker gene expression in the UMAP plots, we used the R package *Nebulosa*^65^ which represents gene expression using kernel density estimation to account for overplotting and noise from expression drop-out.

### Integration of single-cell datasets

Integration of the two scRNA-seq datasets was done using the RPCI method as part of the *RISC* R package^19^, following the suggested pipeline to pre-process raw counts. We used only genes expressed in both datasets to generate 15 gene eigenvectors and using the signalling pathway perturbation dataset as the reference dataset, performed integration to return 50 principal components for further analysis. UMAP dimensionality reduction of the integrated data was also done using the *RISC* package, using 15 components of the ‘PLS’ embeddings as recommended for integrated values. Gene expression UMAP plots were generated using the *Nebulosa* R package. Broad cell type clustering was performed using the *Seurat* package with the resolution parameter set to 0.3.

### Cell-cell interaction analysis

The *CellChat*^21^ R package (v2.1.2) was used to infer, analyse, and visualise cell-cell communication networks. We utilized *CellChatDB* v2, which contains approximately 3,300 validated molecular interactions. In total, we have a combination of 10 treatment conditions (“XAV”, “lowXAV”, “CHIR”, “BMP4”, “Dorso”, “lowDorso”, “K02288”, “VEGF”, “no_treatment”, and “no_treatment_tc”) and three time points (days 2, 5, and 9), resulted in a total of 30 datasets. We applied the CellChat pipeline to each dataset. Briefly, we created the CellChat object based on the corresponding expression matrix and metadata for each dataset with the ‘createCellChat’ function. Communication probability was computed with the ‘computeCommunProb’ function, using the default ‘triMean’ parameter to approximate a 25% truncated mean. This ensures that the average gene expression is set to zero if less than 25% of cells in a group express the gene. The ‘filterCommunication’ function was then applied to filter cell-cell communications using the default setting, requiring a minimum of 10 cells in each group to infer communication. The inferred cellular communication networks were extracted at both the ligand-receptor and signaling pathway levels using the ‘subsetCommunication’ function. Aggregated cell-cell communication networks for each dataset were generated using the ‘aggregateNet’ function. We used CellChat plot functions to visualize three key signaling pathways -WNT, BMP, and VEGF - if they passed statistical significance, along with the top three pathways with the highest probability of cell-cell communication. Network centrality measures from weighted-directed networks were employed to identify the dominant senders, receivers, mediators, and influencers of intercellular communication, using the ‘netAnalysis_computeCentrality’ and ‘netAnalysis_signalingRole_network’ functions. In addition, CellPhoneDB (v5.0.0) along with cellphonedb-data (v5.0), comprising a total of ∼3,000 interactions, was used to infer cell-cell interactions for each of the 30 datasets (**Supplementary File 2**). We generated counts and metadata files from the Seurat object to serve as input for CellphoneDB. Following the guidelines provided in the CellPhoneDB Jupyter notebooks (https://github.com/ventolab/CellphoneDB/tree/master/notebooks), we employed the ‘simple analysis’ mode (method 1) to obtain the mean interaction values for each pair of cell clusters, and employed the ‘statistical_analysis’ mode (method 2) to evaluate the significance of the interactions present in the dataset. Circle plots, including those shown in **Figure S2e**, were generated using the ‘netVisual_aggregate’ function in CellChat with the parameter ‘layout=circle’.

### Gene regulatory network analysis using pySCENIC

To assess regulon activities within individual cells, we used the Python-based package pySCENIC (version 0.12.1)^22,23^. Unlike traditional methods that focus on individual gene expression, pySCENIC determines rank-based regulon activities by inferring co-expression between transcription factor-target modules and identifying enriched cis-regulatory motifs. The analysis began by identifying co-expression modules comprising transcription factors and their potential target genes. This step was performed using the GRNBoost2 algorithm and applied to the gene expression matrix with reference to the hg38 database within pySCENIC. Following the identification of co-expression modules, we performed cis-regulatory motif enrichment analysis to generate candidate transcription factor regulons using cisTarget with the motif collection database version 10 in pySCENIC. Finally, the activity levels of these candidate regulons were quantified at the single-cell level using AUCell. This approach evaluates whether the regulon gene set is significantly enriched at the top of the gene ranking for each cell, providing a measure of regulon activity across different cellular contexts.

### Lineage inference analysis using URD

Differentiation trajectory reconstruction was using the URD method (version 1.1.1)^24^ following the instructions outlined on the authors’ GitHub (https://github.com/farrellja/URD) and utilising default parameters for pre-processing and normalisation of the raw count matrix. For calculation of the diffusion map to determine transition probabilities between cells, 116 nearest neighbours (knn) and a sigma of 20 were used. Next, pseudotime values were assigned to each cell, ordering them along the developmental process. For this, all cells collected from days 2 and 3 of differentiation were used as root cells and 50 flood simulations determined the visitation structure of the lineage specification graph. The URD tree was constructed using all day 9 cells, annotated by their whole-dataset cluster assignment (Figure 2d), as tip cells (terminal cell populations), with a p-value threshold of 0.001, 8 bins per pseudotime window, 25 cells per pseudotime bin, and “preference” as the divergence method.

### TRIAGE-Cluster analysis

To assign each cell a repressive tendency score (RTS), we identified the expressed gene with the highest *TRIAGE* RTS in each cell. The RPCI^19^*-*generated UMAP coordinates were input into the python *scipy*.*stats gaussian_kde* function, using the RTS vector as the “weights” parameter to estimate a probability density function of the repressive tendency across the data in UMAP space. Using a bandwidth of 0.25, a contour representation of the data was generated, splitting the density function into 10 levels. For each level, the spatial clustering algorithm *DBSCAN* from the python module *sklearn* was used to identify all spatially separated groups of cells. *TRIAGE* “peak” regions were found by identifying the *DBSCAN* clusters with no smaller clusters within them at higher contour levels. Code is available at https://github.com/palpant-comp/TRIAGE-Cluster.

### Cell type annotation with reference to *in vivo* datasets

For gene set scoring of cell type domain genes from 921 mouse tissue RNA-seq samples^26^, R package *biomaRt getLDS* function was used to map the mouse Ensembl gene IDs to human Ensembl gene IDs. R package *topGO* was used to determine enriched GO terms for each gene set. For each gene set, the *AddModuleScore* function from the *Seurat* package was used to assign a gene set score for each cell in each cell type peak. After thresholding scores to be greater than 0, the gene set with the highest score was assigned as the cell type domain annotation for that cell. For label transfer, both our query cell type peaks and the reference *in vivo* datasets^2,4^ were *TRIAGE*^25^ transformed as it has been observed to result in higher prediction confidence in label transfer analysis. The label transfer prediction scores were thresholded only to consider cells with scores over 0.5, and the highest scoring cell type was used to annotate each cell. For both gene set scoring and label transfer annotations, each cell peak’s overall annotation is defined as the most common label after thresholding and is assigned as NA if most cells do not have a score greater than the respective thresholds.

### Cell type annotation with reference to spatial domains of embryonic cell types in gastrulating mouse embryo

To annotate early *in vitro* cell types from days 2-4 in our dataset with reference to early *in vivo* cell types, we compared we compared the expression profiles of each *TRIAGE* cell type peak to spatial mouse embryonic transcriptome data that spans embryonic days 5.5 to 7.5^27^. For days 2-4 in our *in vitro* data, the top 50 cell type marker genes were determined for each time point using Cepo, which identifies differentially stable genes between cell types using stability metrics including proportion of zeros and coefficient of variation of a log-transformed normalised expression matrix^67^. Cell type peaks with fewer than 20 cells and genes expressed in less than 5% of any peak were excluded from further analysis. Next, we took the union of all cell type marker genes identified by Cepo and calculated the Pearson correlation of their expression profiles with the spatial mouse transcriptome data. This is used as a proxy to assess how well the expression profile of a cluster recapitulates specific cell types in the gastrulating mouse embryo.

### TRIAGE-ParseR analysis

We calculated the average gene expression value for each *TRIAGE-Cluster* cell type peak and converted these values to the *TRIAGE* discordance score (DS) by multiplication of each gene to its repressive tendency score^25^. We used the top 100 genes ranked by DS from each cell type peak to calculate the pairwise Jaccard similarity index between all peaks. Cell peaks were clustered based on Euclidean distance with complete linkage using the *pheatmap* R package. The top 100 DS genes for each cell peak were then parsed into functionally related modules using gene co-modulation analysis^28^. In brief, principal component (PC) analysis was performed on H3K27me3 deposition breadth data across all genes and cell types in the NIH Roadmap Epigenomics dataset^75^, where each PC represents a biologically meaningful H3K27me3 deposition pattern. The top 67 PCs, which capture over 96% of variance in the data, are used to inform probabilistic clustering of the input genes from each cell peak into co-modulated sets using a Gaussian Mixture Model. The optimal number of clusters is determined by Bayesian Information Criterion, and only clusters with significant (p<0.001, Fisher’s Exact Test) protein-protein interactions as determined by STRING analysis^76^ were kept for further analysis. Code is available at https://github.com/palpant-comp/TRIAGE-ParseR.

### Differential gene expression, KEGG and GO enrichment analysis

For differential gene expression analysis, we utilised the ‘FindMarkerGenes’ function from the *Seurat* R package. Heatmap visualisation displaying number of differentially expressed genes between treatments was generated using the *pheatmap* R package. WNT-related GO term associations were identified using the *biomaRt* R package, and DE genes were annotated by their most specific directional GO term. GO term and KEGG pathway enrichment analysis for the differentially expressed genes compared between the XAV and CHIR treatments at day 9 was performed using the ‘enrichGO’ and ‘enrichKEGG’ functions in the *clusterProfiler* R package. WNT -related genes in Figure 4d are defined as those associated with the following GO terms: Wnt signalling pathway (GO:0016055), Positive regulation of Wnt signalling pathway (GO:0030177), Negative regulation of Wnt signalling pathway (GO:0030178), Positive regulation of canonical Wnt signalling pathway (GO:0090263), Negative regulation of canonical Wnt signalling pathway (GO:0090090), Positive regulation of non-canonical Wnt signalling pathway (GO:2000052), and Negative regulation of non-canonical Wnt signalling pathway (GO:2000051).

### Generation of *TMEM88* KD scRNA-seq dataset

The WTC CRISPRi TMEM88-g2.3 cells in each sample were cultured in 8 temporally staggered setups of mesendoderm-directed differentiation. The ‘XAV’ group was treated with 5uM XAV-393 for 48 hours, from day 3 to day 5 of differentiation, and the ‘TMEM88 KD’ group was treated with 1ug/mL doxycycline for the entire duration of differentiation starting from day 0. Cells were dissociated using 0.5% EDTA in 2.5% Trypsin (ThermoFisher, #15400054) and neutralised with 50% foetal bovine serum (GE Healthcare Life Sciences, #SH30084.03) in DMEM/F12 media (Sigma #11320033). 1 x 10^6^ cells from each sample were labelled with a different TotalSeq-A cell hashing antibody (BioLegend antibodies A2051-8) as per the recommended protocol ^13^ and sorted for viability on a BD Influx Cell Sorter (BD Biosciences) using propidium iodide. 5 x 10^5^ live cells per time point were collected and pooled for Chromium Single Cell 3’ V3 (10x Genomics) reactions following the manufacturer’s protocol, targeting 2 x 10^4^ cells. Gene expression libraries were sequenced on an Illumina NovaSeq 6000 and the cell hashing libraries were sequenced on an Illumina NextSeq 550Dx.

### Quality control processing of *TMEM88* scRNA-seq dataset

This dataset was pre-processed in the same manner as the time course dataset; the cellranger pipeline was used to process raw reads, cell hashing barcodes (time points) were demultiplexed with the ‘HTODemux’ function from the *Seurat* package, and doublet detection was performed with the *scds* package, removing cell barcodes described as doublets by at least 3 methods. Transcriptome-based cell filtering as part of the *Seurat* pipeline removed cells with fewer than 1000 and greater than 10,000 total read counts; fewer than 100 and greater than 300 detected genes; mitochondrial reads comprising greater than 15% of the library; or ribosomal reads making up greater than 45% of the library. Following filtering, time points were assigned to the remaining cells based on their cell hashing barcode with the highest expression in each cell.

### Quantitative RT-PCR

Cells were lysed using β-mercaptoethanol in RLT buffer (QIAGEN) and snap frozen on dry ice. Total RNA was collected with the RNeasy Mini Kit (QIAGEN) according to manufacturer’s instructions. RNA was reverse transcribed into cDNA using Superscript III Reverse Transcriptase (Thermo Fisher Scientific). qRT-PCR was performed using SYBR Green PCR Master Mix reagents on a ViiA 7 Real-Time PCR System (Thermo Fisher Scientific). Gene expression was analysed in GraphPad Prism or R, where relative gene expression levels were calculated for each gene by raising two to the power of the measured gene (averaged over two technical replicates), divided by the mean control housekeeping gene expression. For human samples, *HPRT1* was used as the control (Forward primer 5’-TGA CAC TGG CAA AAC AAT GCA-3’, Reverse primer 5’-GGT CCT TTT CAC CAG CAA GCT-3’), and *Hprt1* was used for mouse samples (Forward primer 5’-TCA GTC AAC GGG GGA CAT AAA-3’, Reverse primer 5’-GGG GCT GTA CTG CTT AAC CAG-3’). Data were analysed in R, where outlier values (those exceeding mean ± 2 standard deviations) in each group were removed. The statistical test used to compare the normalised expression means from separate experiments was a two-tailed *t-*test with a 95% confidence interval using R.

### Flow cytometry

Cells were dissociated with 0.5mM % EDTA + 0.25% Trypsin (1:10) then neutralised with foetal bovine serum (FBS) in DMEM/F12 or RPMI media (1:1). Cells were fixed using 4% paraformaldehyde, permeabilised in 0.75% saponin with 5% FBS in PBS and labelled for flow cytometry using either cardiac troponin T (Creative Biolabs, #M40283), KDR (R&D Systems, #FAB357P), or CD34 (BD Biosciences, #340430) with the corresponding immunoglobin G isotype control. Flow cytometry data was captured using a BD FACSCANTOII with the FACDiva software. Analysis of this data was performed using *FlowJo* (v10.07). Summaries of the cell populations were analysed and visualised in R, where outlier values (exceeding mean ± 2 standard deviations) in each group were removed before performing a two-tailed *t*-test for the comparison of means with a 95% confidence interval.

### Summary-data based Mendelian Randomisation (SMR) analysis

We used 61 anthropometric and cardiovascular phenotypes ^32^ from the UK Biobank^77^, whose heritability estimates were given as “medium” or “high” by Neale’s lab, to explore the broad effect of select genes on human traits. Publicly available summary statistics generated by Neale’s lab were used to perform Summary-data Mendelian Randomisation (SMR) analysis using eQTL (expression quantitative trail loci) information derived in the V8 GTEX release^78^. Individuals with European ancestry from the 1000 genomes project were used as a LD (linkage disequilibrium) reference. SMR analysis was performed with a p-value threshold for eQTL inclusion of 0.05 to ensure all eQTLs affecting gene expression were captured. SMR analysis across all phenotypes was performed on the vascular (coronary artery, tibial artery, aortic artery), cardiac (left ventricle, atrial appendage), respiratory (lung), and gut tube derivative (gastroesophageal junction, oesophageal mucosa, pancreas, stomach, and thyroid) tissues. Further analysis for *TMEM88* was performed on systolic blood pressure (4080) and diastolic blood pressure (4079) phenotypes using all 49 GTEX tissues.

### Generation of *Tmem88* conditional knockout mouse line

The floxed *Tmem88* mice were generated by the Monash Genome Modification Platform, Monash University (see acknowledgements). In brief, CRISPR RNAs were ordered from Integrated DNA Technologies and annealed with trans-activating CRISPR RNA (Alt-R CRISPR-Cas9 tracrRNA from IDT #107319 to form single guide RNA (gRNA) targeting Exon 2 of Tmem88 (gRNA1: 5’-AAT GGG CCA CTC TGC CCA GG-3’, gRNA2: 5’-TTT GCC CAT TTG TAG TCT TG-3’). 10ng/uL Cas9 nuclease (IDT Alt-R® S.p. Cas9 nuclease) was incubated with 10ng/uL gRNAs to form a ribonucleoprotein (RNP) complex. A 1247bp ssDNA repair template containing LoxP sites flanking the second exon of *Tmem88* (see also Figure 5i) was generated using Invitrogen Dynabeads™ MyOne™ Streptavidin C1 according to manufacturer’s instructions. The RNP and ssDNA repair template (10ng/uL) were microinjected into the pronucleus of C57BL/6J zygotes at the pronuclei stage. Microinjected zygotes were transferred into the uterus of pseudo pregnant F1 females.

All mouse breeding was conducted in accordance with consent from the University of Queensland’s Molecular Biosciences Animal Ethics committee approval (Molecular Biosciences AEC-MBS approval #: 2018/AE000177 & 2021/AE000421). Floxed *Tmem88* animals were crossed with C57BL/6 mice expressing Cre recombinase driven by a CMV promoter, to generate animals with ubiquitous loss of the *Tmem88* gene. Both hetero- and homozygous mutants were viable and showed no obvious phenotype. For genotyping the animals, DNA was extracted from ear or toe samples using 50uL of QuickExtract (Biosearch Technologies, #QE09050), incubated at 65°C for 15 minutes, then 100°C for 5 mins and diluted to 50ng/uL for genotyping. Each mouse was tested for genomic presence of Cre-Recombinase (Forward primer 5’-CTG ACC GTA CAC CAA AAT TTG CCT G-3’, Reverse primer 5’-GAT AAT CGC GAA CAT CTT CAG GTT C-3’; Cre = 200bp), Floxed *Tmem88* (Forward primer 5’-CGG CTC AGA ATC TGC TGG TG-3’, Reverse Primer 5’-TTT CTA GCG GGA AGC GGT GT-3’; Wt = 1300bp, Flox = 1350bp, KO = 600bp), and both LoxP sites (5’LoxP Forward primer 5’-CGG CTC AGA ATC TGC TGG TG-3’, 5’LoxP Reverse primer 5’-GCA GAT GCG AGG TGC AAA GG-3’, 3’LoxP Forward primer 5’-TCC AAT CGC TCC TGC ACT GG-3’, 3’LoxP Reverse primer 5’-TTT CTA GCG GGA AGC GGT GT-3’; Wt = 300bp, LoxP = 350bp). Quantitative RT-PCR was also used to confirm the loss of Tmem88 expression in the mutants (Forward primer 5’-CTG GTG GCT GTT TTC AAT CTC C-3’, Reverse primer 5’-GTG CCT GAG AGC GCA GAA A-3’). For RNA extraction, 30mg heart tissue was homogenised in liquid nitrogen using a mortar and pestle before proceeding with the RNeasy Mini Kit (QIAGEN #74106) following the manufacturer’s instructions. cDNA synthesis and quantitative RT-PCR were performed as described above.

### Mouse echocardiography

All mouse experiments were carried out in accordance with consent from the University of Queensland’s Molecular Biosciences Animal Ethics committee approval (Molecular Biosciences AEC-MBS approval #: 2018/AE000171 & 2021/AE000999). Blinded echocardiography was performed on 8 week old mice using a Vevo 3100 Preclinial Imaging System (VisualSonics, Toronto, Canada) with a 25-55 MHz linear transducer (MX550D). Mice were anaesthetised using 4% isofluorane, and general anaesthesia was maintained with 1% isofluorane during the procedure. Mice were weighed and placed in supine position on a heating pad during echocardiography with body temperature, heart rate, and electrocardiography recorded throughout. Warmed echo transmission gel was applied to the hairless chest of the mouse before collecting images. Two-dimensional B-mode images recorded in the parasternal long axis view were used to determine left ventricular end-diastolic volume (LVEDV). One-dimensional M-mode images in the parasternal short-axis view were also used to measure ejection fraction (EF) and thicknesses of the posterior walls of the left ventricle in diastole (LVPWd). Pulsed-wave doppler recordings in apical four-chamber view were used to measure mitral inflow velocity of blood during passive filling of the ventricle (E-wave) and active filling during atrial systole (A-wave). Pulsed-wave tissue doppler in apical four-chamber view was used to capture mitral annular velocity during early (E’) and late (A’) diastole. Analysis of echocardiography images was performed in VevoLab (version 3.1.1) software (VisualSonics, Toronto, Canada). LVEDV, and LVPWd were normalised to body weight to achieve left ventricle end diastolic index (LVEDI/LVESI), and LVPWd index, respectively. For each animal, all parameters were measured on average 4 times (minimum twice) and averages are presented. Animals with abnormal body weights and heart rates were excluded (4 animals removed), as well as all measurements from images with consistent outlier measurements indicating incorrect image capture (12 images removed). Statistical analysis was carried out using R with the *ggpubr* package to perform a two-sided *t-*test with a 95% confidence interval.

### Mouse blood pressure measurements

Adult mice between 8 and 28 weeks of age were anaesthetised using 4% isofluorane before endotracheal intubation and mechanical ventilation. Isofluorane was reduced to 1.5% to maintain general anaesthesia for the remainder of the procedure. A Millar Pressure-Volume (PV) catheter (Millar #SPR-839, 1.4F Catheter) was used to cannulate the right carotid artery, to allow a stable pressure trace as measured by the MVPS Ultra Pressure-Volume Unit (Millar, #880-0168). The continuous baseline blood pressure was measured for 5-10 minutes and recorded using the LabChart software (ADInstruments, Dunedin, New Zealand). The average maximum (systolic) and minimum (diastolic) blood pressure across the pressure cycles in the measured time frame are presented.

### Micro-computed tomography and automated image analysis (LAMA)

Embryos were dissected at embryonic day (E)15.5 in PBS, rinsed in cold water and stained in 0.7% phosphotungstic acid diluted in 70% ethanol at RT for 14 days^79^. Stained embryos were mounted in agarose and scanned on a Skyscan 1272 micro-CT scanner (Bruker) using an Al 0.5 + Cu 0.038 filter and voltage of 87 kV. Images were captured at 1640 × 2452 resolution (6 μm^3^ voxel size) through 360° at 0.4° angle increments and reconstructed with smoothing, ring artefact reduction and beam hardening options in NRecon software (Bruker). 3D datasets were downsampled to 14 μm^3^ and each registered using the LAMA phenotyping pipeline^80^ to an E15.5 population average so that its 43-organ atlas (in preparation) maps onto each embryo dataset. The resulting organ and whole embryo volumes (WEV) were fitted to a linear model organ volume/WEV∼genotype+WEV with heterozygous and homozygous *Tmem88* embryos statistically compared to wildtype littermate controls. P-values were corrected for multiple comparisons using the LAMA permutation statistics script.

## MATERIALS AVAILABILITY

Details for all resources presented in this study are available at our web accessible dashboard (http://cellfateexplorer.d24h.hk/) and will be made available upon request.

## DATA AND CODE AVAILABILITY

All three single-cell RNA sequencing raw datasets generated in this study are available upon request. Additionally, all figures, data, and an interactive version of the combined, annotated dataset are available at http://cellfateexplorer.d24h.hk/, where additional co-modulation analysis outputs, metadata, and gene expression for individual cell type peaks can be queried. *TRIAGE-Cluster* and *TRIAGE-ParseR* scripts are available at https://github.com/palpant-comp/TRIAGE-Cluster and https://github.com/palpant-comp/TRIAGE-ParseR. Raw and processed sequencing data will be made available on the Gene Expression Omnibus platform.

