## Supplemental Figures, Supplemental Tables 1-2 for "A pluripotent stem cell atlas of multilineage differentiation reveals *TMEM88* as a developmental regulator of mammalian blood pressure"

#### SUPPLEMENTARY FIGURES AND TABLES

**Figure S1**: Quality control and barcoding metrics of scRNA-seq datasets.

**Figure S2**: Characterisation of atlas dataset with broad cell type clusters.

**Figure S3**: Characterisation of atlas dataset with *TRIAGE-Cluster* cell peaks.

**Figure S4**: Evaluation of WNT signalling and candidate genes regulating differentiation.

**Figure S5**: *TMEM88* as a regulator of mesendodermal development in hiPSCs and mouse model.

**Table S1**: Quality control and barcoding metrics in iPSC pilot scRNA-seq experiment.

**Table S2**: Quality control and barcoding metrics in atlas of differentiation dataset.

**Table S3**: All CellChat significant ligand-receptor interactions.

**Table S4**: Enrichment of all regulons' activity in each cluster from pySCENIC analysis.

**Table S5**: *TRIAGE* genes used for identifying each cell type peak in the atlas dataset.

**Table S6**: Top 100 *TRIAGE* identity-defining genes for each *TRIAGE-Cluster* cell type peak and their *TRIAGE-ParseR* gene cluster assignments in the atlas dataset.

**Table S7**: Specificity of cell identity-defining *TRIAGE-ParseR* genes across all *TRIAGE-Cluster* cell type peaks in the atlas dataset. .

**Table S8**: GO terms enriched for each *TRIAGE-ParseR* gene cluster in each *TRIAGE-Cluster* cell type peak in the atlas dataset.

**Table S9**: SMR analysis results for the top 100 most enriched genes as well as all differentially Wnt-related genes' association to systolic and diastolic blood pressure in cardiovascular tissue.

**Supplementary File 1**: Barcoding plasmid DNA sequence.

**Supplementary File 2**: CellphoneDB cell-cell interaction enrichment tables.

### SUPPLEMENTARY FIGURES

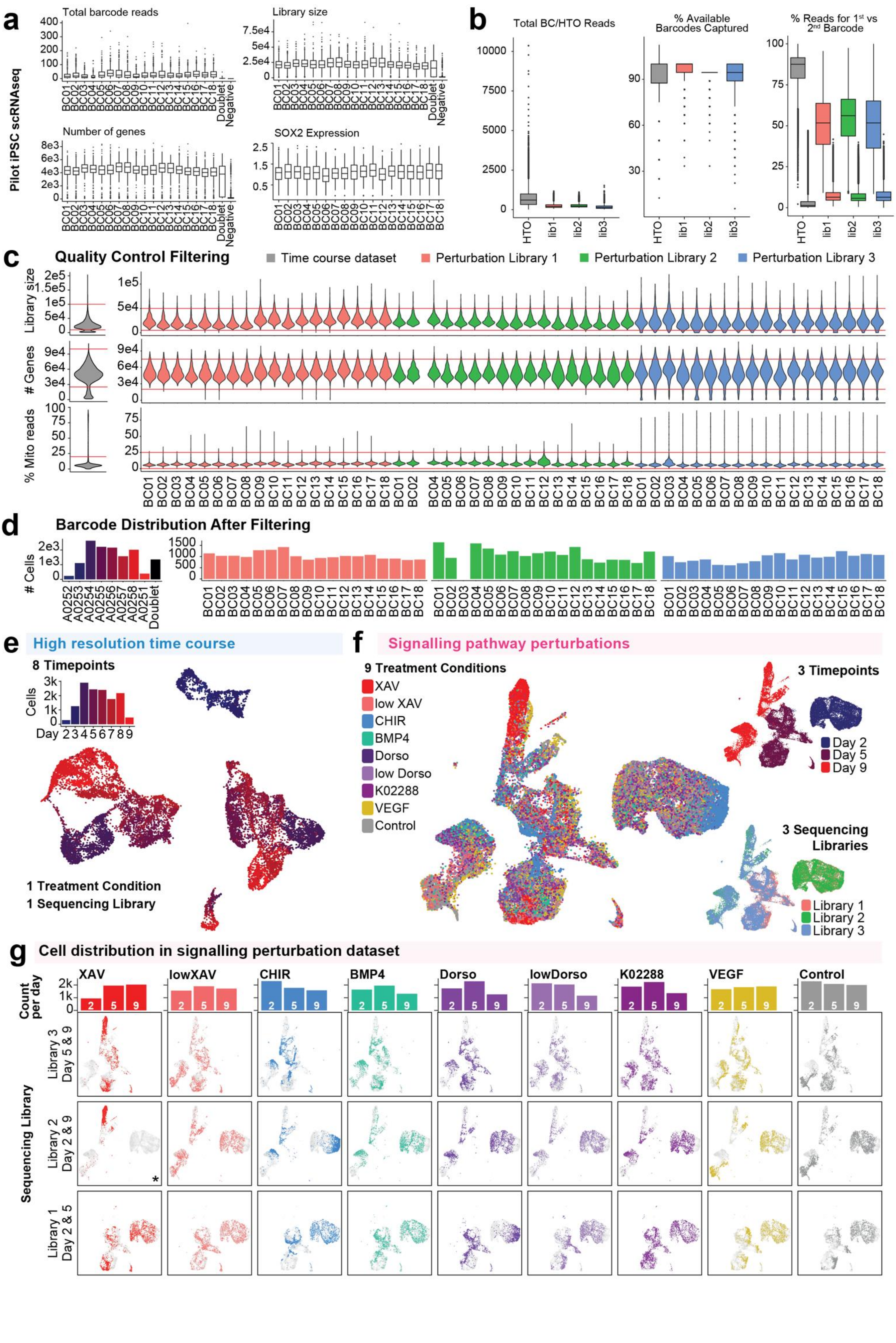

**Figure S1: Quality control of engineered barcoding cell lines.**

**a)** Box plots showing total number of genomic barcode reads (top left); total number of gene expression reads (library size, top right); total number of captured genes (bottom left); and expression of pluripotency gene *SOX2* in the pilot iPSC scRNA-seq experiment, grouped by the most highly mapped barcode per cell. See also **Table S1**.
**b)** Box plots showing total cell hashing (HTO) and genomic barcode (BC/libs 1-3) reads (left), percentage of total possible hashing or genomic barcodes captured (middle), and percentage of hashing or genomic barcode reads mapping to the two most highly mapped barcode per cell (right). See also **Table S2**.
**c)** Violin plots showing distribution of three quality control metrics and filtering thresholds chosen, compared between each cell line and scRNA-seq library.
**d)** Distribution of cells after demultiplexing to allocate cells to each of the eight Cell Hashing antibodies, doublet, and negative cell barcode identities after filtering the cells for quality control metrics and reassignment of doublets not supported by multiple doublet detection algorithms (right). Bar plots showing number of cells annotated with an internal barcode versus doublet or negative classifications after re-allocation doublet cells not supported by multiple doublet detection algorithms.
**e)** UMAP plot showing all high quality cells (13,682 cells total) in the time course data after filtering. Cells are coloured by time point.

**f)** UMAP plot showing all high quality cells (48,526 cells total) in the signalling perturbation dataset after standard filtering and processing. Cells are coloured by treatment (upper), time point (lower left), and sequencing library (lower right). See **Table 1** for detailed breakdown of treatment, timepoint, and sequencing library combinations used.

**g)** Distribution of cells from each treatment, timepoint, and sequencing library in the signalling perturbation dataset. * NB: one of the day 2 replicates in the XAV group was missing from the experiment (see also **Table 1**, **Figure S1c & d**).

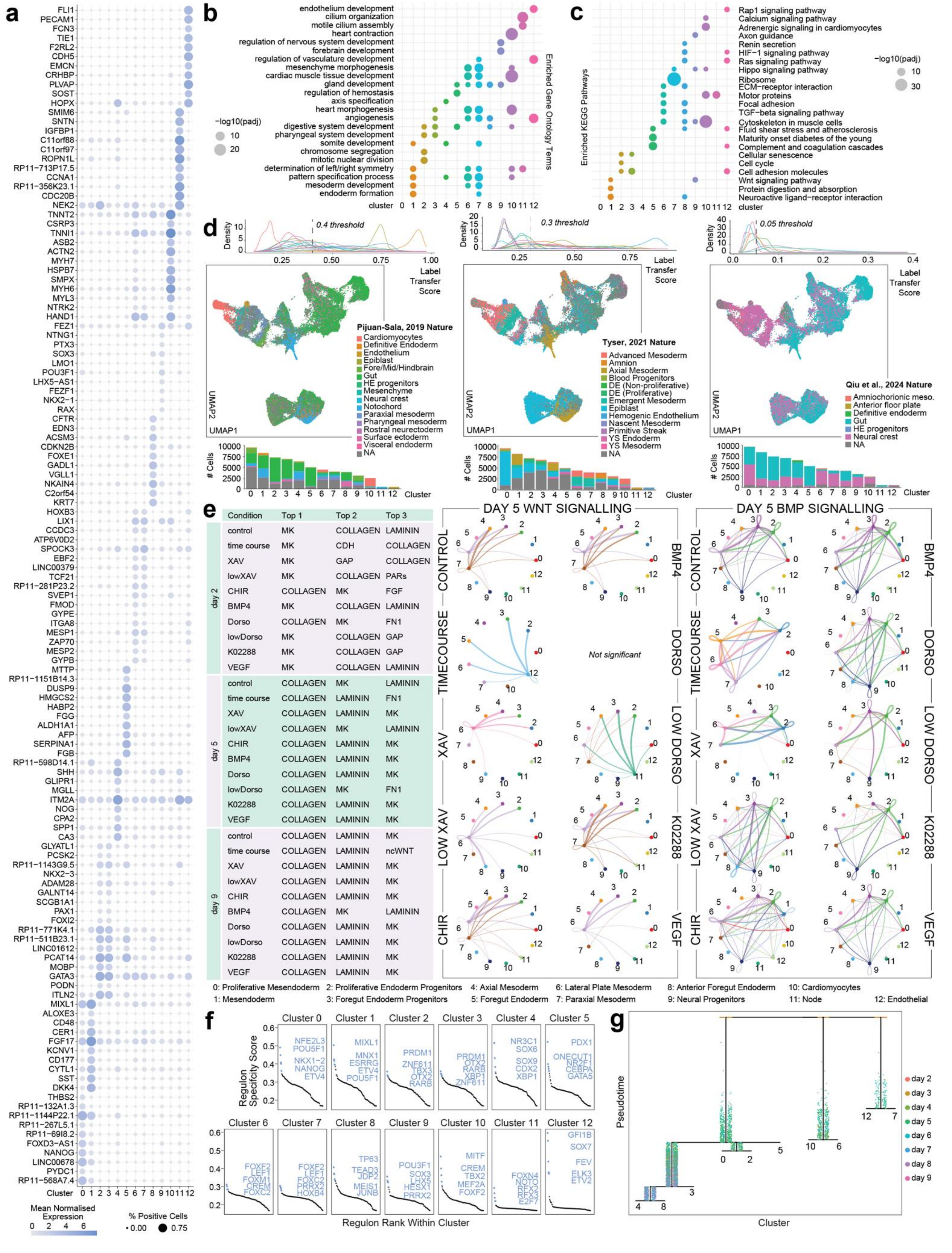


| | Condition | Top 1 | Top 2 | Top 3 |
|---|---|---|---|---|
| day 2 | control | MK | COLLAGEN | LAMININ |
| | time course | MK | CDH | COLLAGEN |
| | XAV | MK | GAP | COLLAGEN |
| | lowXAV | MK | COLLAGEN | PARs |
| | CHIR | COLLAGEN | MK | FGF |
| | BMP4 | MK | COLLAGEN | LAMININ |
| | Dorso | COLLAGEN | MK | FN1 |
| | lowDorso | MK | COLLAGEN | GAP |
| | K02288 | MK | COLLAGEN | GAP |
| | VEGF | MK | COLLAGEN | LAMININ |
| day 5 | control | COLLAGEN | MK | LAMININ |
| | time course | COLLAGEN | LAMININ | FN1 |
| | XAV | COLLAGEN | LAMININ | MK |
| | lowXAV | COLLAGEN | MK | LAMININ |
| | CHIR | COLLAGEN | LAMININ | MK |
| | BMP4 | COLLAGEN | LAMININ | MK |
| | Dorso | COLLAGEN | LAMININ | MK |
| | lowDorso | COLLAGEN | MK | FN1 |
| | K02288 | COLLAGEN | LAMININ | MK |
| | VEGF | COLLAGEN | LAMININ | MK |
| day 9 | control | COLLAGEN | LAMININ | MK |
| | time course | COLLAGEN | LAMININ | ncWNT |
| | XAV | COLLAGEN | LAMININ | MK |
| | lowXAV | COLLAGEN | LAMININ | MK |
| | CHIR | COLLAGEN | LAMININ | MK |
| | BMP4 | COLLAGEN | MK | LAMININ |
| | Dorso | COLLAGEN | LAMININ | MK |
| | lowDorso | COLLAGEN | LAMININ | MK |
| | K02288 | COLLAGEN | LAMININ | MK |
| | VEGF | COLLAGEN | LAMININ | MK |

**Figure S2: Characterisation of atlas dataset with broad cell type clusters.**

**a)** Expression of marker genes for each cell cluster. Included genes were selected from known marker genes as well as the top 10 differentially expressed genes for each cluster.

**b-c)** Enrichment of select GO terms (**b**) and KEGG pathways (**c**) based on the top 100 most significantly differentially expressed genes for each cluster. Relevant terms and pathways were selected from the top 10 most significantly enriched results for each cluster.
**d)** Label transfer results assessing similarity of each single-cell against *in vivo* cell types in mouse (left[1], right[2]) and human (middle[3]) embryogenesis. For each reference dataset we show distribution of label transfer prediction scores are shown with the selected threshold (top), a UMAP with cells with scores above the threshold coloured by their most confidence label (middle), and proportion of cells from each cluster annotated with each reference label (bottom). HE: Haematoendothelial; NA: Cells with scores below the threshold cut-off; DE: Definitive Endoderm; YS: Yolk Sac; Amniochorionic meso.: Amnichorionic mesoderm B.
**e)** Results from CellChat predicting cell-cell interactions compared between treatment groups. The 3 pathways with the highest probabilities for each group at days 2, 5, and 9 of differentiation are displayed in the table (left), and significant predicted WNT- (centre) and BMP- (right) pathway communication between cell type clusters on day 5 are provided. See also **Table S3** for full CellChat output of all predicted ligand-receptor interactions between clusters at all time points.
**f)** Significantly enriched gene regulon activity in each cell cluster based on identification transcription factor-target co-expression modules and cis-regulatory motif enrichment analysis as part of the pySCENIC pipeline. Regulons are ranked by their specificity score (y-axis) and the top 5 most enriched regulons are highlighted. See also **Table S4** for the whole enrichment output for all regulons in each cluster.
**g)** Reconstruction of differentiation trajectories using the URD algorithm[4]. Clusters present at day 9 of differentiation were used as tips for tree construction.

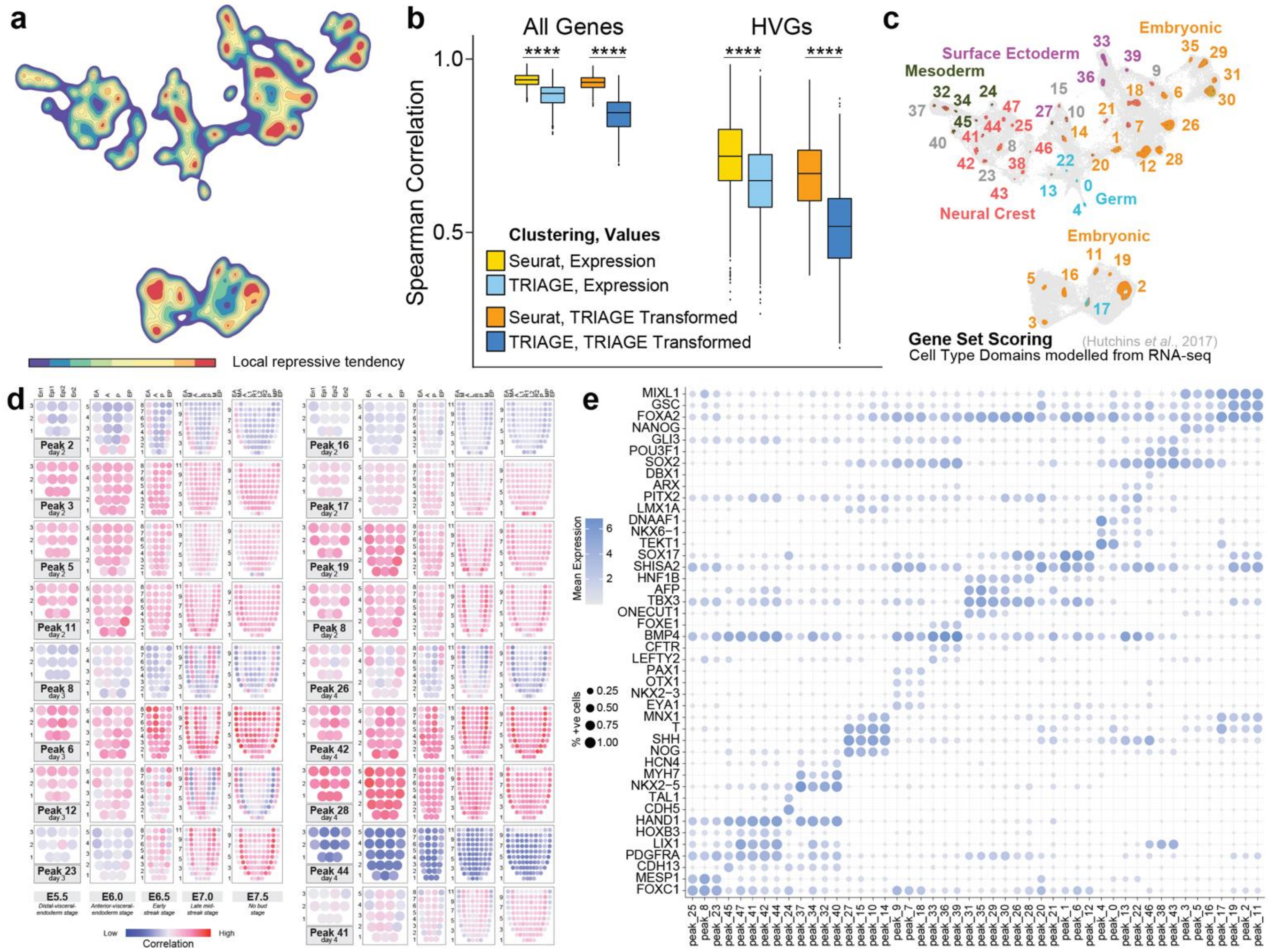


**Figure S3: Quality control and filtering of scRNA-seq experiments.**

**a)** Contour plot showing regions of cells in the UMAP space expressing genes with high repressive tendency scores using bandwidth 0.25 to inform *TRIAGE-Cluster*[5] analysis.

**b)** Comparison of peak-to-peak transcriptomic similarity (Spearman rank correlation) using all genes (left) and just the top 1000 highly variable genes (right), compared between Seurat clustering (resolution 2.1 to achieve the same number of clusters identified using *TRIAGE-Cluster*) and *TRIAGE-Cluster*, before and after *TRIAGE* transformation[6] of the expression values. ****$p<2.22e-16$.

**c)** Annotation of *TRIAGE* cell type peaks by reference to broad cell type domains from gene sets from 921 mouse tissue RNA-seq samples[7] using *Seurat* gene set scoring. Cells are coloured on a single-cell basis by their assigned label after thresholding (threshold = 0). Dark grey cells do not have a label above threshold. Peak number labels are coloured by the label that makes up the greatest proportion of cells in that peak, where grey labels indicate peaks where most cells do not score above the threshold. Light grey cells do not fall within a peak. See also **Figure 3b-c**.

**d)** Annotation of early (days 2-4) *TRIAGE* cell type peaks with reference to spatial gene expression domains in the developing mouse embryo at five different stages of development between embryonic days 5.5 and 7.5, spanning pre-gastrulation to late primitive streak and amnion formation[8,9]. For each peak and timepoint, the five corn plots capture different stages of development in mouse embryos, where each spot has captured the expression profiles of approximately 5-40 cells. En1 and En2: divided endoderm; Epi1 and Epi2: divided epiblast; EA: anterior endoderm; A: anterior; P: posterior; EP: posterior endoderm; M: whole mesoderm; L: left lateral; R: right lateral; MA: anterior mesoderm; L1: anterior left lateral; R1: anterior right lateral; L2: posterior left lateral; R2: posterior right lateral; MP: posterior mesoderm. See Figure 3k for expert annotation of highly correlated domains for each peak and timepoint.

**e)** Expression of select marker genes for each *TRIAGE* cell type peak to support annotations in **Figure 3h**.

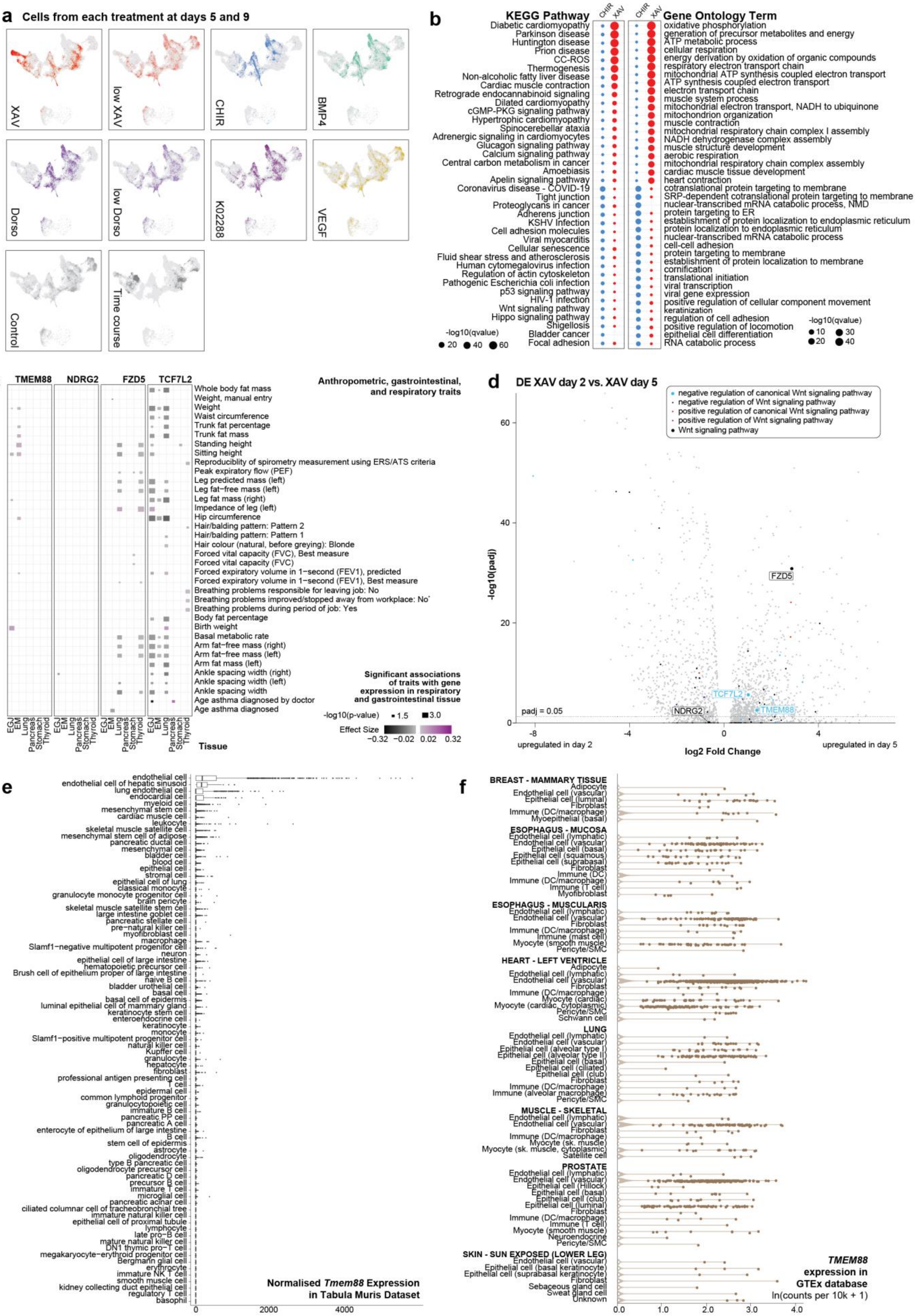


**Figure S4: Evaluation of Wnt signalling and candidate genes regulating differentiation.**

**a)** Proportion of cells from each treatment condition allocated to each cell type peak at days 5 and 9 of differentiation.

**b)** Enrichment of KEGG pathways and GO terms in genes that are differentially expressed between XAV and CHIR-treated cells at day 9. CC-ROS: chemical carcinogenesis-reactive oxygen species; KSHV: Kaposi sarcoma-associated herpesvirus; HIV: Human immunodeficiency virus; NMD: nonsense-mediated decay.

**c)** SMR analysis evaluating associations of candidate WNT-related gene expression in respiratory or gastrointestinal tissues with anthropometric, gastrointestinal, and respiratory traits in the UK Biobank. Some trait and tissue names were shortened from the following: *Breathing problems improved/stopped away from workplace or on holiday: No; EGJ: Esophagus gastroesophageal junction; EM: Esophagus mucosa.

**d)** Differential gene expression analysis comparing XAV-treated cells at days 2 and 5 of differentiation. Genes annotated with the listed GO terms related to the Wnt signalling pathway are coloured and the four candidate Wnt regulators from **Figure 4d-e** are labelled.

**e)** *Tmem88* gene expression from Tabula Muris dataset[10].

**f)** *TMEM88* gene expression from GTEx[11].

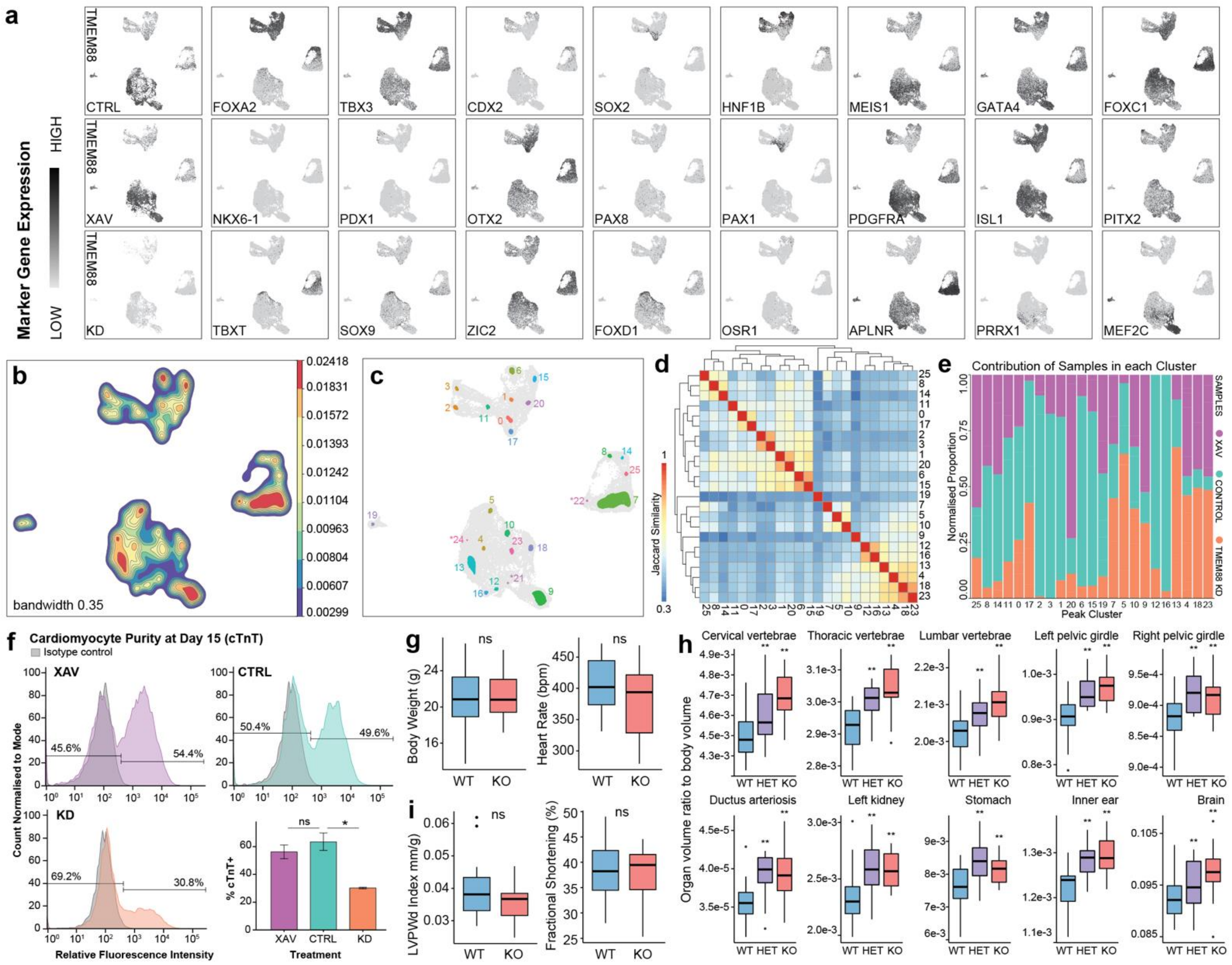

**Figure S5: *TMEM88* as a regulator of mesendodermal development in hiPSCs and mouse model.**

**a)** Expression of additional cell type marker genes in the *TMEM88* scRNA-seq dataset. First column shows expression of *TMEM88* compared between the three treatment conditions. *FOXA2* and *TBX3* mark definitive endoderm, *CDX2* marks hindgut, *SOX2* marks pluripotency and anterior foregut, *HNF1B* marks liver progenitors, *MEIS1* and *GATA4* mark mesoderm, *FOXC1* marks somitic mesoderm, *NKX6-1* and *PDX1* mark pancreatic progenitors, *OTX2* marks anterior foregut, *PAX8* marks thyroid progenitors, *PAX1* marks pharyngeal endoderm, *PDGFRA* and *ISL1* mark lateral plate mesoderm, *PITX2* and *PRRX1* mark limb bud mesoderm, *T* and *SOX9* mark axial mesoderm, *ZIC2* and *FOXD1* mark neuromesodermal neural crest progenitors, *OSR1* marks intermediate mesoderm, *ALPNR* marks lateral plate mesoderm, and *MEF2C* marks cardiac mesoderm.

**b)** Contour plot showing regions of cells in the UMAP space expressing genes with high repressive tendency scores using bandwidth 0.35 to inform *TRIAGE-Cluster*[5] analysis.

**c)** UMAP showing immediate output from *TRIAGE-Cluster*, identifying 26 distinct cell types. Cell type labels with an * have fewer than 20 cells and are removed in further analysis.

**d)** Heatmap and dendrogram showing pairwise Jaccard similarity between the top 100 genes for each peak after *TRIAGE*-transformation[6].

**e)** Contribution of cells from each sample to each cell type peak in the dataset. Within each column, proportions sum to 1 and the input number of cells for each sample is normalised to the total number of cells collected in that sample.

**f)** Representative plots from FACS analysis for cTnT+ (cardiac troponin T) cells from day 15 of mesendoderm differentiation (left) and summarised FACS results (n = 2-4, right). Data are represented as mean ± standard error. * $p \leq 0.05$, ns = not significant, as determined by a Wilcoxon signed rank test.

**g)** Comparison of overall body weight and heart rates between wildtype and *Tmem88* KO mice.

**h)** Summary of micro-computed tomography results showing significant difference in normalised organ volume between heterozygous or homozygous *Tmem88* mutant E15.5 embryos and wildtype littermates. P-values were adjusted for multiple comparisons by permutation-based correction to an FDR of 0.01. ** $p<0.01$.

**i)** Echocardiographic analysis of heart function in *Tmem88* KO vs control. Left ventricular posterior wall at diastole (LVPWd) index normalised to body weight. n=13-22, * $p < 0.05$ by *t*-test.

#### SUPPLEMENTARY TABLES

**Table S1:** Quality control and barcoding metrics in iPSC pilot scRNA-seq experiment.

| Group | Highest BC/HTO | # cells | % high QC cells | Median total RNA reads | Median total genes captured | Median #BC/HTO reads | Median # mapped BC/HTOs | Median 1st BC/HTO Count | Median 2nd BC/HTO Count |
|---|---|---|---|---|---|---|---|---|---|
| All barcoded cells | BC01 | 421 | 91.21 | 21032 | 4365 | 16 | 2 | 15 | 1 |
| | BC02 | 473 | 89.22 | 19924 | 4254 | 19 | 1 | 18 | 0 |
| | BC03 | 231 | 92.64 | 23127 | 4707 | 13 | 2 | 12 | 1 |
| | BC04 | 265 | 90.94 | 22652 | 4540 | 10 | 2 | 9 | 1 |
| | BC05 | 226 | 88.05 | 21561 | 4442 | 23 | 2 | 23 | 1 |
| | BC06 | 315 | 85.40 | 21130 | 4379 | 37 | 2 | 36 | 1 |
| | BC07 | 226 | 79.65 | 24494 | 4922 | 25 | 1 | 24 | 0 |
| | BC08 | 222 | 79.28 | 25161 | 4888 | 23 | 2 | 22 | 1 |
| | BC09 | 188 | 93.09 | 21355 | 4395 | 14 | 2 | 14 | 1 |
| | BC10 | 184 | 88.59 | 21054 | 4432 | 20 | 2 | 19 | 1 |
| | BC11 | 381 | 87.14 | 21669 | 4391 | 22 | 2 | 21 | 1 |
| | BC12 | 188 | 83.51 | 24205 | 4721 | 23 | 2 | 22 | 1 |
| | BC13 | 186 | 82.26 | 24114 | 4690 | 20 | 2 | 18 | 1 |
| | BC14 | 325 | 92.31 | 21185 | 4423 | 19 | 2 | 18 | 1 |
| | BC15 | 448 | 90.85 | 19942 | 4184 | 25 | 2 | 23 | 1 |
| | BC16 | 339 | 92.33 | 20310 | 4190 | 30 | 1 | 29 | 0 |
| | BC17 | 379 | 89.71 | 18206 | 4039 | 26 | 2 | 25 | 1 |
| | BC18 | 332 | 88.25 | 19199 | 4121 | 28 | 1 | 27 | 0 |
| | No BC | 1299 | 4.93 | 1017 | 119 | 0 | 0 | 0 | 0 |
| | Doublet* | 83 | 56.63 | 15696 | 3796 | 2 | 2 | 1 | 1 |
| | All cells | 6711 | 71.96 | 18944 | 4121 | 15 | 1 | 14 | 0 |
| All hashed cells | HTO-A2051 (BC07) | 209 | 85.17 | 24059 | 4773 | 1 | 1 | 1 | 0 |
| | HTO-A2052 (BC08) | 160 | 91.25 | 25880 | 4913 | 1 | 1 | 1 | 0 |
| | HTO-A2053 (BC12) | 251 | 88.84 | 23105 | 4610 | 1 | 1 | 1 | 0 |
| | HTO-A2054 (BC13) | 259 | 91.89 | 23697 | 4701 | 1 | 1 | 1 | 0 |
| | HTO Doublet* | 45 | 100.00 | 30328 | 5217 | 2 | 2 | 1 | 1 |
| | All BC + HTO | 924 | 89.83 | 24066 | 4733 | 1 | 1 | 1 | 0 |
| Barcoded cells negative for hashing oligos | BC07 (No HTO) | 93 | 79.57 | 22983 | 4814 | 24 | 1 | 22 | 0 |
| | BC08 (No HTO) | 141 | 70.92 | 22470 | 4658 | 19 | 1 | 18 | 0 |
| | BC12 (No HTO) | 78 | 76.92 | 23259 | 4632 | 19 | 1 | 18 | 0 |
| | BC13 (No HTO) | 95 | 76.84 | 21018 | 4426 | 17 | 2 | 16 | 1 |
| | BC07,08,12,13 (No HTO) | 407 | 75.43 | 22367 | 4623 | 19 | 1 | 18 | 0 |
| Hashed cells negative for barcodes | HTO-A2051 (No BC) | 68 | 4.41 | 1312 | 108 | 1 | 1 | 1 | 0 |
| | HTO-A2052 (No BC) | 41 | 2.44 | 1151 | 113 | 1 | 1 | 1 | 0 |
| | HTO-A2053 (No BC) | 45 | 2.22 | 1066 | 106 | 1 | 1 | 1 | 0 |
| | HTO-A2054 (No BC) | 25 | 8.00 | 1273 | 148 | 1 | 1 | 1 | 0 |
| | HTO Doublet (No BC)* | 1 | 0.00 | 1279 | 216 | 1 | 2 | 1 | 1 |
| | All HTO only (no BC) | 180 | 3.89 | 1227 | 113 | 2 | 1 | 1 | 0 |

*Doublet cells refer to cells with exactly the same number of reads mapped to >1 barcode or HTO.

**Table S2:** Quality control and barcoding metrics in atlas of differentiation dataset.

| Sample/Library | Cell Count (Pre-filter) | Median Library Size | Median # Genes | Median % Mt | Median % Rb | Median BC/HTO reads | Median BC/HTOs captured | Median %reads for 1st BC/HTO | Median %reads for 2nd BC/HTO | % Negative BC/HTO | % Doublet BC/HTO | % Singlet BC/HTO | Final Cell Count (post QC) |
|---|---|---|---|---|---|---|---|---|---|---|---|---|---|
| **Time Course (HTO)** | 19997 | 23575 | 4984 | 5.83 | 27.93 | 605 | 7 | 87.53 | 1.36 | 1.75 | 20.09 | 78.17 | 13682 |
| **Signalling Lib 1 (BC)** | 19656 | 22812 | 4926 | 6.89 | 29.15 | 204 | 18 | 51.77 | 6.44 | 2.17 | 23.83 | 74.00 | 16586 |
| **Signalling Lib 2 (BC)** | 19708 | 21801 | 4761 | 8.06 | 27.29 | 225 | 17 | 56.14 | 5.73 | 0.13 | 22.42 | 77.45 | 16391 |
| **Signalling Lib 3 (BC)** | 19349 | 21063 | 4992 | 5.86 | 21.63 | 142 | 17 | 51.75 | 6.40 | 10.42 | 23.83 | 65.75 | 15549 |
| **Control Day 2 (HTO-A0252)** | 531 | 23483 | 4768 | 8.38 | 28.74 | 541 | 8 | 88.25 | 1.44 | 0.19 | 20.53 | 79.28 | 330 |
| **Control Day 2 (Lib 2, BC01)** | 1664 | 22108 | 4517.5 | 8.14 | 33.86 | 215 | 17 | 56.26 | 5.64 | 0.06 | 15.02 | 84.92 | 1635 |
| **Control Day 2 (Lib 1, BC10)** | 1003 | 28483 | 5264 | 7.72 | 32.01 | 248 | 18 | 52.90 | 6.15 | 0.90 | 31.21 | 67.90 | 934 |
| **Control Day 5 (HTO-A0255)** | 3192 | 18284.5 | 4228 | 3.57 | 34.34 | 315 | 7 | 79.68 | 2.17 | 8.49 | 12.91 | 78.60 | 2687 |
| **Control Day 5 (Lib 1, BC01)** | 1203 | 22137 | 4906 | 5.57 | 29.77 | 223 | 18 | 58.91 | 5.41 | 2.08 | 21.11 | 76.81 | 1142 |
| **Control Day 5 (Lib 3, BC10)** | 1304 | 22574 | 5184 | 6.01 | 22.42 | 168.5 | 18 | 56.67 | 5.59 | 7.29 | 25.54 | 67.18 | 1148 |
| **Control Day 9 (HTO-A0251)** | 744 | 21076.5 | 4674.5 | 5.61 | 25.07 | 887 | 8 | 91.99 | 1.10 | 0.13 | 27.82 | 72.04 | 605 |
| **Control Day 9 (Lib 3, BC01)** | 1165 | 22433 | 4913 | 5.47 | 25.20 | 162 | 18 | 55.89 | 5.77 | 7.38 | 26.27 | 66.35 | 1017 |
| **Control Day 9 (Lib2, BC10)** | 1265 | 19625 | 4705 | 6.85 | 24.66 | 222 | 17 | 60.18 | 5.37 | 0.16 | 23.79 | 76.05 | 1218 |

See separate .xlsx files for **Tables S3-9**.
